# Experience-dependent, sexually dimorphic synaptic connectivity defined by sex-specific cadherin expression

**DOI:** 10.1101/2024.05.08.593207

**Authors:** Chien-Po Liao, Maryam Majeed, Oliver Hobert

## Abstract

We describe here the molecular mechanisms by which juvenile experience defines patterns of sexually dimorphic synaptic connectivity in the adult nervous system of the nematode *C. elegans*. We show that starvation of juvenile males disrupts serotonin-dependent activation of the CREB transcription factor in a nociceptive sensory neuron, PHB. CREB acts through a cascade of transcription factors to control expression of an atypical cadherin protein, FMI-1/Flamingo. During postembryonic development, FMI-1/Flamingo has the capacity to promote and maintain synaptic connectivity of the PHB nociceptive sensory to a command interneuron, AVA, in both sexes, but the serotonin transcriptional regulatory cassette antagonizes FMI-1/Flamingo expression in males, thereby establishing sexually dimorphic connectivity between PHB and AVA. A critical regulatory node in this process is the CREB-target LIN-29, a Zn finger transcription factor which integrates four different layers of information – sexual specificity, past feeding status, time and cell-type specificity. Our findings provide the mechanistic details of how an early juvenile experience defines sexually dimorphic synaptic connectivity.

## INTRODUCTION

Early life experiences, such as stress and depression, can affect later developmental processes and can also impact on the manifestation of neurological disorders (LUPIEN *et al*. 2009; HERRINGA *et al*. 2013). Notably, these past experiences can generate outcomes in a sex-dependent manner. For example, clinical studies indicate that females have a higher chance of developing anxiety or depression when experiencing early-life adversity (HISCOX *et al*. 2023). However, how past experiences influence later aspects of nervous system function in a sex-dependent manner remains elusive in humans. In rodents, a recent study has shown that juvenile adversity alters corticolimbic connectivity in traumatized female rat but not the male counterpart, and consequently, female rats are more likely to display depression behavior (HONEYCUTT *et al*. 2020). Here again, the molecular mechanisms that drive such sexually dimorphic outcomes are unknown.

Using the nematode *C. elegans* as a model, we have previously shown that the experience of early juvenile starvation affects the establishment of sexually dimorphic synaptic connectivity during sexual maturation (BAYER AND HOBERT 2018). Specifically, the phasmid sensory neuron PHB generates *en passant* synapses onto the command interneuron AVA in both sexes at early juvenile stages; however, upon sexual maturation this synaptic connection is sex-specifically maintained and progressively strengthened in hermaphrodites but eliminated in males (WHITE *et al*. 1986; JARRELL *et al*. 2012; OREN-SUISSA *et al*. 2016; BAYER AND HOBERT 2018; COOK *et al*. 2019). Early starvation increases the expression of the invertebrate norepinephrine analog octopamine to inhibit serotonin release from a pair of head sensory neurons (BAYER AND HOBERT 2018). This transient serotonin depletion and consequent lack of activation of the metabotropic SER-4 receptor in PHB, results in the failure of the PHB>AVA synaptic pruning and instead promotes growth of PHB>AVA synaptic connectivity in males, thereby eliminating sexually dimorphic connectivity (BAYER AND HOBERT 2018).

Serotonin-mediated activation of metabotropic receptors can affect a wide range of signaling pathways (BOCKAERT *et al*. 2006). We show here that the relevant read-out in sex-specific serotonin-mediated synaptic pruning is a multilayered transcriptional response, in which serotonin signaling first activates the CREB transcription factors. CREB then directly activates the Zn finger transcription factor LIN-29A, a key regulatory node in this process.

Transcription of the *lin-29a* locus bookmarks feeding status via CREB activation, which cooperates with a cell-type specific terminal selector to direct *lin-29* expression to specific neuron types. Activation of *lin-29a* transcription is antagonized in hermaphrodites by the TRA-1 master regulatory of sexual identity. *lin-29a* transcripts are translationally inhibited by the LIN-41 RNA binding protein until sexual maturation. Once LIN-29A protein is produced at the right place and time, it directs sexually dimorphic PHB>AVA connectivity by repression of another transcription factor, the Doublesex transcription factor, DMD-4, which we had previously found to be expressed in PHB of hermaphrodites, but not males (BAYER *et al*. 2020a). We identify the non-conventional cadherin *fmi-1/Flamingo* as the terminal effector of the CREB>LIN-29A>DMD-4 transcription factor cascade. We show that FMI-1 is normally required in hermaphrodites to promote the increase in the number of PHB>AVA *en passant* synapses. Male-specific repression of FMI-1 via serotonin/CREB-mediated LIN-29A induction (and DMD-4 repression) therefore leads to a failure to sustain and expand PHB>AVA synapse number. Hence, we have discovered a mechanism whereby food perception in early life is translated into the control of the later development of a sexually dimorphic synaptic connectivity.

## RESULTS

### Visualization of sexually dimorphic PHB>AVA connectivity and neurite contact length and its differential dependence on past experience

Juvenile starvation (starving animals at the L1 stage for 24 hours and transferring them back to food) affects sexually dimorphic PHB>AVA synaptic number decrease at the later sexual maturation stage via temporally mediating serotonin (5-HT) signaling in the sensory neuron PHB (BAYER AND HOBERT 2018). The molecular pathway underlying such experience-dependent, sex-specific synaptogenesis remains unknown. We had previously visualized the effect of feeding state and 5-HT signaling on PHB>AVA synaptic connectivity through the use of “GFP-reconstitution across synaptic partner” (GRASP) technology which exploits the synaptically localized neuroligin protein NLG-1 (FEINBERG *et al*. 2008; BAYER AND HOBERT 2018). We validated and expanded our previous results through the establishment and usage of additional reagents. First, an independently generated PHB>AVA GRASP reporter transgene, *otIs839* confirmed our previous results in experience-dependent sexual dimorphic connectivity of PHB and AVA (BAYER AND HOBERT 2018)(**Figure 1A,B**). Second, we confirmed that NLG-1-based GRASP is indeed a proper indicator of PHB>AVA synaptic connectivity by using a GFP-tagged synaptic active zone marker, CLA-1/Clarinet, expressed specifically in PHB, as well as a postsynaptic marker, the ionotropic glutamate receptor AVR- 14 which is expressed in the AVA neurons, the postsynaptic target of the glutamatergic PHB neurons (GAT *et al*. 2023; LI *et al*. 2023). While GFP::CLA-1 alone labels all synaptic outputs of PHB, including those to many male-specific neurons, adjacent localization of PHB- expressed GFP::CLA-1 and AVA-expressed AVR-14::TagRFP signals represents an indicator of PHB>AVA connectivity (**Figure 1C-F**). We found that either in isolation or in combination, the number of GFP::CLA-1 and AVR-14::TagRFP signals corroborate the conclusions based on the GRASP constructs: The number of *en passant* PHB>AVA synaptic signals show (a) no dimorphisms in juvenile stages, (b) display dimorphisms in after sexual maturation and (c) starvation results in aberrant *en passant* synapse number increase in males (**Figure 1E-F, Figure S1A,B**).

**Figure 1.**
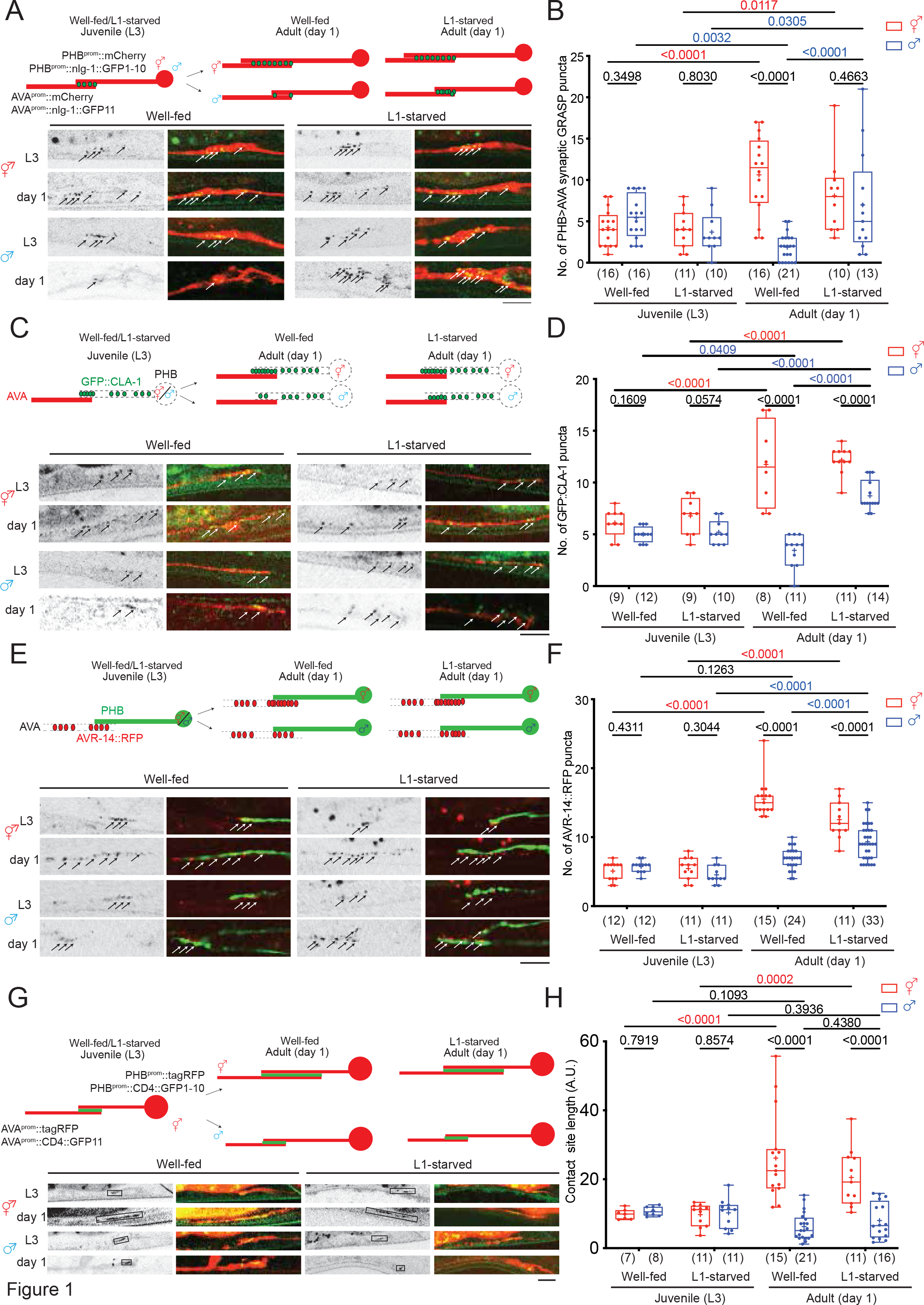
Juvenile serotonin signaling patterns sexually dimorphic synaptic connectivity. **(A,B)** Representative images (A) and quantification (B) of PHB>AVA synaptic GRASP(*otIs839*) in L3 and day 1 well-fed and L1-starved animals in both sexes. **(C)**(top) Schematic illustration of and (bottom) Representative images of AVA-juxtaposed GFP::CLA-1 in PHB (*otIs883;otEx8040*) in L3 and day 1 well-fed and L1-starved animals in both sexes. **(D)** Quantification of AVA-juxtaposed GFP::CLA-1 in PHB in L3 and day 1 well-fed and L1- starved animals in both sexes. Note for panel C and D that the number of AVA-juxtaposed CLA-1 puncta remained sexually dimorphic in the L1-starved adult male, which is consistent with electron micrographic data that shows that PHB generates many sex-specific synapses, in addition to AVA (COOK *et al*. 2019). **(E)** (top) Schematic diagram and (bottom) representative images of PHB-juxtaposed AVR- 14::TagRFP in in L3 and day 1 well-fed and L1-starved animals in both sexes. **(F)** Quantification of AVR-14::TagRFP in AVA (*otIs902; him-8(e1489)*) in L3 and day 1 well- fed and L1-starved animals in both sexes. **(G)** (top) Schematic diagram and (bottom) representative images of PHB>AVA neurite CD4- GRASP (*otEx8152*) in L3 and day 1 well-fed and L1-starved animals in both sexes. We measure the GFP-positive length to indicate the PHB/AVA contact site. **(H)** Quantification of CD4-GRASP (*otEx8152*) in L3 and day 1 well-fed and L1-starved animals in both sexes. Statistics: (B,D,F,H) Two-way ANOVA followed by Bonferroni multiple comparisons test. *p*- value and N numbers are indicated on the graph. + indicates the mean value. Scale bar = 10 µm.

Our previous global analysis of neurite adjacency and *en passant* synapse formation throughout the *C. elegans* nervous system indicates that the extent of adjacency of two neurites is a sufficient predictor of the number of synapses formed between adjacent neurons (COOK *et al*. 2023). Since the PHB>AVA synapses are generated *en passant* along the PHB and AVA neurites, we considered the possibility that the extent of neurite adjacency is also sexually dimorphic and regulated by juvenile experience. We visualized PHB/AVA neurite contact length by using the transmembrane CD4 protein to direct the two halves of GFP (GFP1-10 and GFP11) to the surface of the PHB and AVA membranes, respectively. As previously demonstrated in other *C. elegans* cell types (FEINBERG *et al*. 2008), GFP should only reconstitute when the two neurite membranes contact each other. We measured GFP- positive neurite length to indicate the length of the PHB/AVA contact site **(Figure 1G)**. We found that in juvenile animals, the PHB/AVA contact length was comparable between both sexes; however, in day 1 adults, the contact length was significantly increased in hermaphrodites but not in males **(Figure 1G, H)**. This altered neurite contact length is confirmed by simply examining cytoplasmic reporters that fill AVA and PHB axons (data not shown), but due to the limits of resolution, it is only through the use of split GFP technology, that we can confirm that such adjacency is in the molecular range. Unexpectedly, while juvenile starvation affects the manifestation of sexually dimorphic synapse number, it did not affect sexually dimorphic neurite contact length **(Figure 1G, H).** Hence, the extent of sexually dimorphic neurite contact and sexually dimorphic synapse can be uncoupled. Below, we define a molecular pathway that is dedicated toward controlling sexually dimorphic synaptogenesis, without affecting sexually dimorphic extent of neurite contact.

### CRH-1/CREB functions downstream of juvenile serotonin signals to control male-specific synaptic remodeling upon sexual maturation

We have previously shown that juvenile starvation operates via the disruption of serotonin signaling through the metabotropic serotonin receptor SER-4, to then control PHB>AVA *en passant* synapse number growth and elimination (BAYER AND HOBERT 2018). To dissect the serotonin-triggered signaling cascade in the PHB neuron, we expressed specifically in the PHB neurons a gain-of-function version (gof) of the G-alpha protein, GOA- 1, that we hypothesized to act downstream of the SER-4 G-protein-coupled serotonin receptor (GUREL *et al*. 2012). We found that the loss of PHB>AVA synaptic dimorphisms in adult males that were starved at the L1 stage were rescued in such transgenic animals **(Figure 2A)**, indicating that GOA-1 signaling in male PHB neurons promotes the male-specific diminishment of PHB>AVA *en passant* synapses. This result is further substantiated by genetic removal of the 5-HT synthesizing enzyme, TPH-1. In *tph-1* mutant males, sexually dimorphic PHB>AVA connectivity was disrupted, based on GRASP, CLA-1 and AVR-14 punctae **(Figure 2B**, **S1C, D)** and these defects were rescued by PHB-specific GOA-1^gof^ expression **(Figure 2C)**.

**Figure 2.**
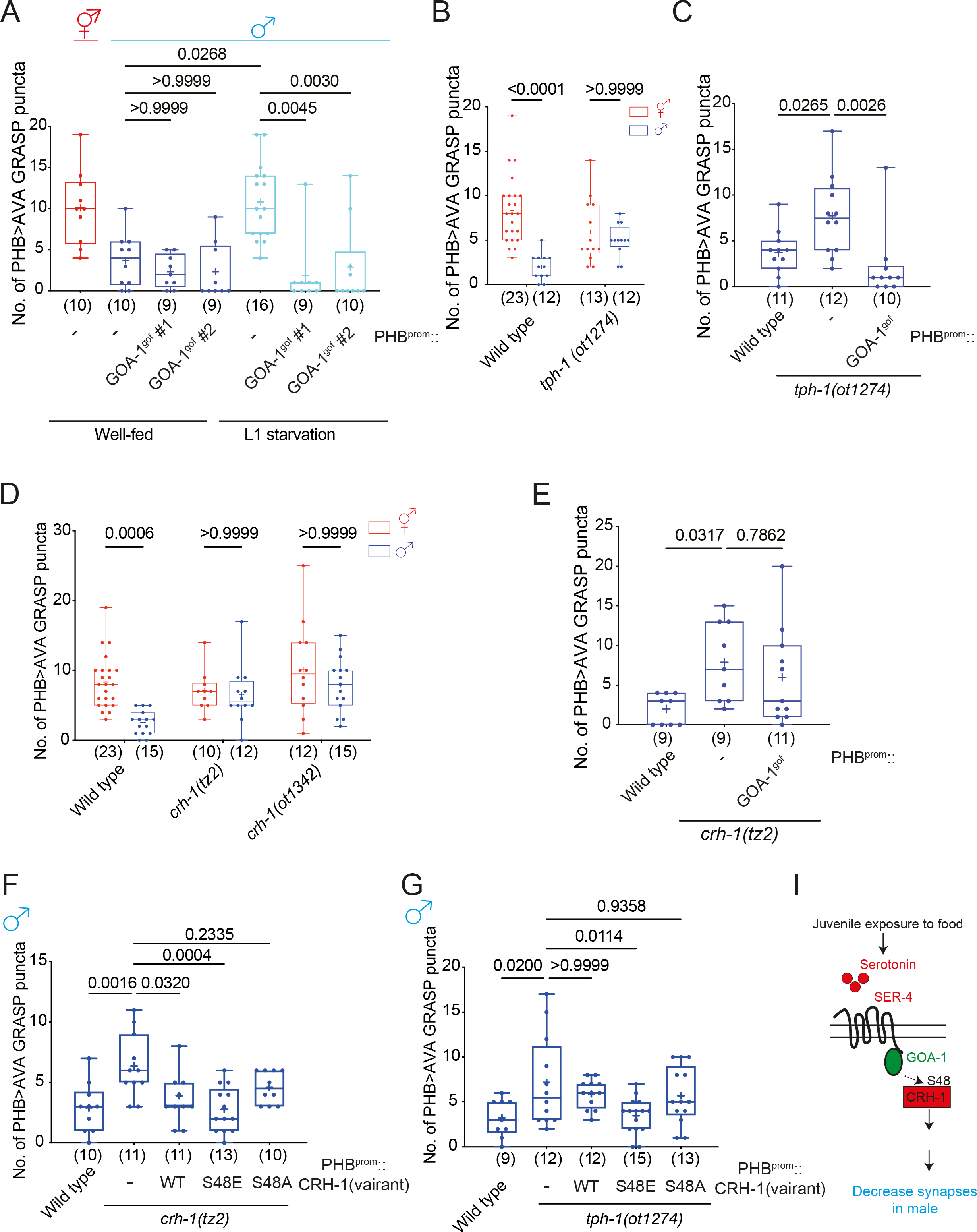
Juvenile starvation controls sexually dimorphic PHB>AVA synaptic contacts via CRH-1/CREB. **(A)** Quantification of PHB>AVA synaptic GRASP(*otIs839*) in well-fed wild-type animals and L1-starved wild-type and animals expressing PHB::LIN-29A (*otEx7915* ) or PHB::GOA- 1^gof^(*otEx7925* and *otEx8158*). **(B)** Quantification of PHB>AVA synaptic GRASP (*otIs839*) in wild-type and *tph-1(ot1274)* in both sexes. **(C)** Quantification of PHB>AVA synaptic GRASP (*otIs839*) in wild-type and *tph-1(ot1274)* males expressing PHB::GOA-1^gof^(*otEx7925* and *otEx8158*). **(D)** Quantification of PHB>AVA synaptic GRAPS (*otIs839*) in wild-type and *crh-1(tz2)* and *crh-1(ot1342)* animals of both sexes. **(E)** Quantification of PHB>AVA synaptic GRASP (*otIs839*) in wild-type and *crh-1(tz2)* males expressing PHB::GOA-1^gof^(*otEx7925* and *otEx8158*). **(F,G)** Quantification of PHB>AVA synaptic GRASP(*otIs839*) in *crh-1(tz2)* (F) and *tph- 1(ot1274)* (G) males with overexpressing CRH-1 missense allele transgenes (*otEx8045* for CRH-1^WT^, *otEx8046* for CRH-1^S48E^, and *otEx8082* for CRH-1^S48A^) in the PHB neurons. **(I)** Schematic diagram indicates juvenile food experience acts through serotonin-GPCR-GOA- 1 to activate CRH-1 to secure LIN-29A expression upon sexual maturation to establish PHB>AVA sexually dimorphic connectivity. Statistics: (A,C,E,F,G) One-way ANOVA and (B,D,I) two-way ANOVA followed by Bonferroni multiple comparisons test. *p*-value and N numbers are indicated on the graph. Scale bar = 10 µm. + indicates the mean value.

Among the many downstream effector pathways of metabotropic 5HT signaling is the activation of CREB via protein phosphorylation (LONZE AND GINTY 2002; BOCKAERT *et al*. 2006; OURY *et al*. 2010; ZHANG *et al*. 2016). We analyzed two independent *crh-1* loss of function alleles, *tz2* and *ot1342,* and found that PHB>AVA sexually dimorphic synaptic patterning was lost in these animals **(Figure 2D).** PHB-specific GOA-1^gof^ failed to rescue ectopic PHB>AVA synaptic defects in *crh-1* mutants **(Figure 2E)**, confirming that CRH-1 functions downstream of 5-HT-GPCR signaling to mediate adult male specific PHB>AVA remodeling. We also made use of the fact that CREB proteins, including *C. elegans* CRH-1, are activated by upstream G-protein signaling via a defined phosphorylation site, serine 48 (S48) in *C. elegans* (S133 in mammalian CREB)(KIMURA *et al*. 2002; LONZE AND GINTY 2002). A phosphorylation-deficient mutation (S48A) is predicted to inactivate the protein, while a phosphomimetic mutation (S48E) is predicted to make CRH-1 independent of an upstream- activating input (in this case, loss of serotonin signaling). Indeed, we found that restoration of wild-type and CRH-1^S48E^ but not CRH-1^S48A^ in the PHB rescued PHB>AVA defects in the *crh- 1* mutant males **(Figure 2F)**. Moreover, only restoration of phosphomimetic CRH-1^S48E^ but not wild-type or CRH-1^S48A^ into the PHB neuron rescued the PHB>AVA synaptic defects of serotonin-deficient *tph-1* mutant males **(Figure 2G)**, consistent with CRH-1 acting downstream of serotonin **(Figure 2I)**.

### CRH-1/CREB controls feeding state-dependent male-specific LIN-29A expression

We identified a functionally relevant transcriptional target of CRH-1/CREB by turning to the Zn finger transcription factor LIN-29A, which we had previously shown to be expressed in several neuron classes, including PHB and AVA, only in males, but not hermaphrodites (PEREIRA *et al*. 2019). We found that juvenile experience impacts proper LIN-29A expression in sexually mature males by starving animals at the L1 stage, transferring them back to food, and examining the expression of a CRISPR/Cas9-engineered reporter allele of *lin-29a*. We observed that 80% of the animals show an obvious decrease (reduction or complete elimination) in LIN-29A protein expression (**Figure 3A, B**). We supplemented well-fed animals with octopamine and found that such treatment reduced LIN-29A expression, hence recapitulating the effect of starvation (**Figure S2A**). Corroborating the previously reported critical window period at which starvation affects PHB>AVA synaptic remodeling (**Figure S1E**)(BAYER AND HOBERT 2018), we observed that octopamine exposure at L1 but not the L3 stage results in reduced LIN-29A protein expression (**Figure S2A**). On the other hand, exogenously supplying 5-HT during L1-starvation rescued the LIN-29A expression deficiency (**Figure S2A**). This result is further confirmed by the demonstration that genetic removal of endogenous 5-HT by using animals that lack the 5-HT synthesizing enzyme, TPH-1, diminishes LIN-29A expression in PHB **(Figure 3C).** PHB-specific GOA^gof^ overexpression rescued the L1-starvation LIN-29A expression defects **(Figure 3D)**.

**Figure 3.**
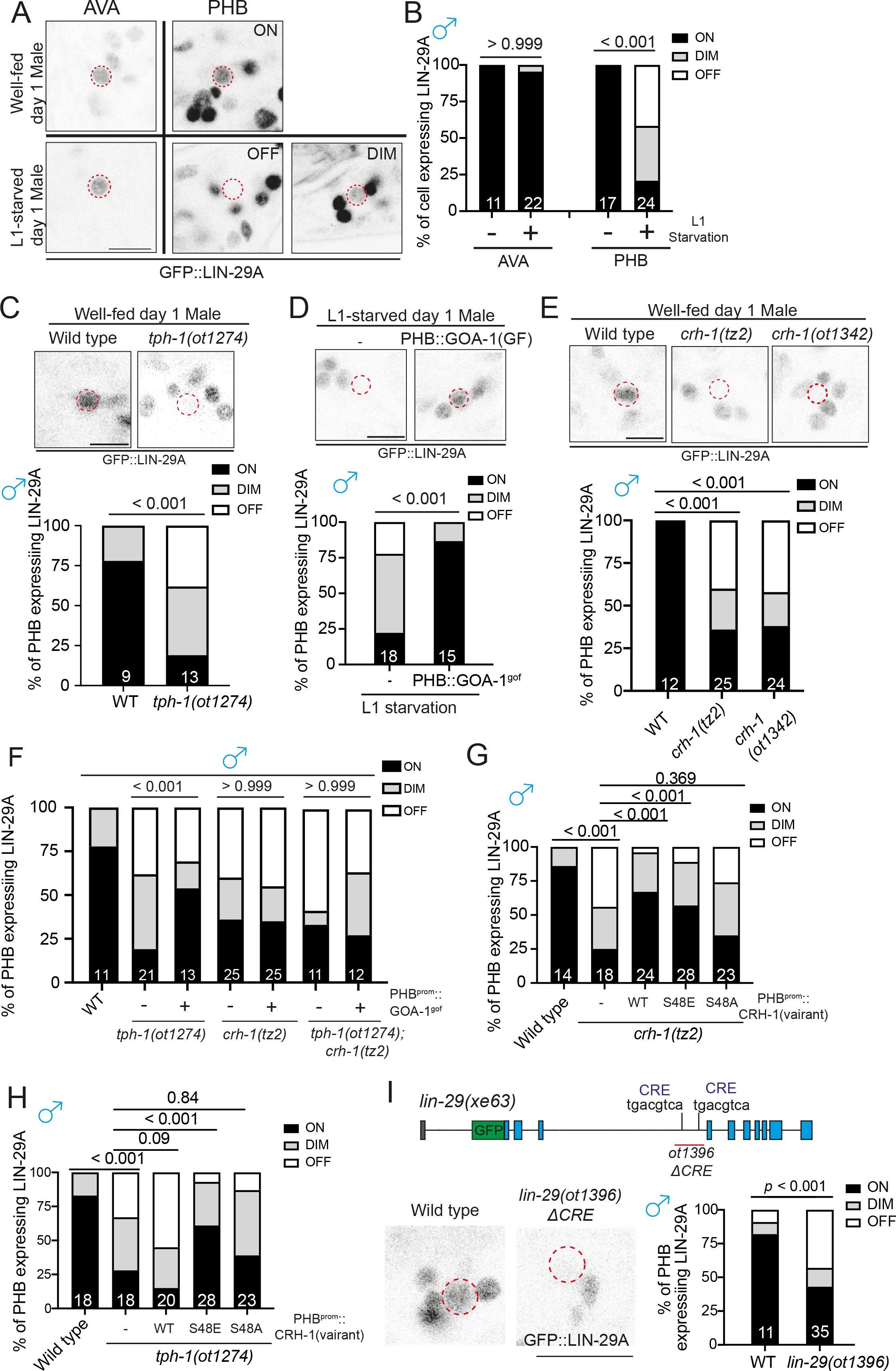
Male-specific LIN-29A expression in PHB is controlled by juvenile serotonin experience via CRH-1/CREB. **(A)** *lin-29(xe63[gfp::lin-29a])* expression of well-fed and L1-starved day 1 males in the AVA and PHB neurons. *lin-29(xe63[gfp::lin-29a])* expression is not affected in the AVA but is dim or lost in the PHB neurons when males undergo L1 starvation. **(B)** Quantification of the percentage of neurons expressing *lin-29(xe63[gfp::lin-29a])* in AVA or PHB under well-fed or L1 starvation conditions. **(C)** Representative images (top) and quantification of (bottom) PHB neurons expressing *lin- 29(xe63[gfp::lin-29a])* in wild-type and *tph-1(ot1274)*. **(D)** Representative images (top) and quantification of (bottom) PHB neurons expressing *lin- 29(xe63[gfp::lin-29a])* in males that undergo L1-starvation with transgene overexpressing GOA-1^gof^ (*otEx8037*) in the PHB neurons. **(E)** Representative images (top) and quantification of (bottom) PHB neurons expressing *lin- 29(xe63[gfp::lin-29a])* in wild-type, *crh-1(tz2)* and *crh-1(ot1342)*. **(F)** Quantification of PHB neurons expressing *lin-29(xe63[gfp::lin-29a])* in animals with or without PHB::GOA-1^gof^ transgene (*otEx8037*) overexpression in wild-type, *tph-1(ot1274)*, *crh- 1(tz2)* and *tph-1(1274); crh-1(tz2)* background. **(G)** Quantification of PHB neurons expressing *lin-29(xe63[gfp::lin-29a])* in *crh-1(tz2)* mutants with overexpressing CRH-1 missense allele transgenes (*otEx8053* for CRH-1^WT^, *otEx8157* for CRH-1^S48E^, and *otEx8113* for CRH-1^S48A^) in the PHB neurons. **(H)** Quantification of PHB neurons expressing *lin-29(xe63[gfp::lin-29a])* in *tph-1(ot1274)* mutants, overexpressing distinct types of *crh-1* transgenes (*otEx8053* for CRH-1^WT^, *otEx8157* for CRH-1^S48E^, and *otEx8113* for CRH-1^S48A^) in the PHB neurons. **(I)** (top) Schematic illustration of “<u>C</u>REB <u>R</u>esponsive <u>E</u>lement” (CRE) sites in the *lin-29a* locus. The *lin-29(ot1396)* allele is designed to delete the potential CRE sites at the intron 3 of the *lin-29a* locus. (bottom) Representative images (left) and quantification of (right) PHB neurons expressing GFP::LIN-29A in wild-type and *lin-29(ot1396)*. Statistics: *chi*-squared tests followed by Bonferroni multiple comparisons test. *p*-value and N numbers are indicated on the graph. The red dashed circle indicates PHB. Scale bar = 5 µm.

Male-specific LIN-29A expression in PHB was also diminished in two independent *crh- 1* alleles **(Figure 3E)**. PHB-specific GOA^gof^ overexpression restored LIN-29A expression defects in *tph-1* mutants but not in *crh-1* or *tph-1; crh-1* double mutant males, consistent with GOA-1 acting downstream of serotonin, but upstream of CRH-1 activation **(Figure 3F).**

Restoration of wild-type and CRH-1^S48E^ but not CRH-1^S48A^ CRH-1 rescued LIN-29A expression defects in the *crh-1* mutant males (**Figure 3G**). Moreover, only restoration of phosphomimetic CRH-1^S48E^ but not wild-type or CRH-1^S48A^ rescued the reduction of LIN-29A expression in serotonin-deficient *tph-1* mutant males (**Figure 3H**). The effect of CRH-1/CREB in LIN-29A expression is likely to be direct, since deletion of putative “<u>C</u>REB <u>R</u>esponsive <u>E</u>lements” (CRE) in the third intron of the *lin-29a* gene also phenocopied defective LIN-29A expression in *crh-1* mutants (**Figure 3I**).

### Sexual, spatial and temporal specificity of LIN-29A expression

The feeding state-dependent control of LIN-29A protein appearance in the male PHB neurons illustrates a fascinatingly complex regulation of LIN-29A protein expression and, importantly, raises the question of how a signal perceived at the L1 stage is translated into male-specific LIN-29A protein appearance at later larval stages. We further investigated all axes of LIN-29 regulation, i.e. its cell-type/spatial specificity (PHB neuron), its sexual specificity (in males), its temporal specificity (protein occurrence during sexual maturation) and feeding state-dependence. We find that the cell-type specificity of induction of LIN-29A in PHB requires the terminal selector of PHB identity, the *ceh-14* LIM homeobox gene (KAGOSHIMA *et al*. 2013; SERRANO-SAIZ *et al*. 2013)(**Figure 4A**). The sexual specificity of LIN- 29A induction in male PHB and not hermaphrodite PHB is, in turn, specified by the global master regulator of sexual identity, the Zn finger transcription factor TRA-1, since removal of TRA-1 selectively in PHB results in LIN-29A depression in hermaphrodite PHB (PEREIRA *et al*. 2019)(**Figure 4B**). CREB activation cannot overcome TRA-1-dependent, sex-specific repression since the PHB::GOA-1^gof^ transgene is insufficient to induce ectopic LIN-29A protein expression in hermaphrodites (**Figure 4C**). We, therefore, surmise that the activity of CREB is directed to *lin-29a* by the presence of neuron-type specific cofactors, CEH-14, and this activation is antagonized in hermaphrodites by the master regulator of hermaphroditic sex, TRA-1.

**Figure 4.**
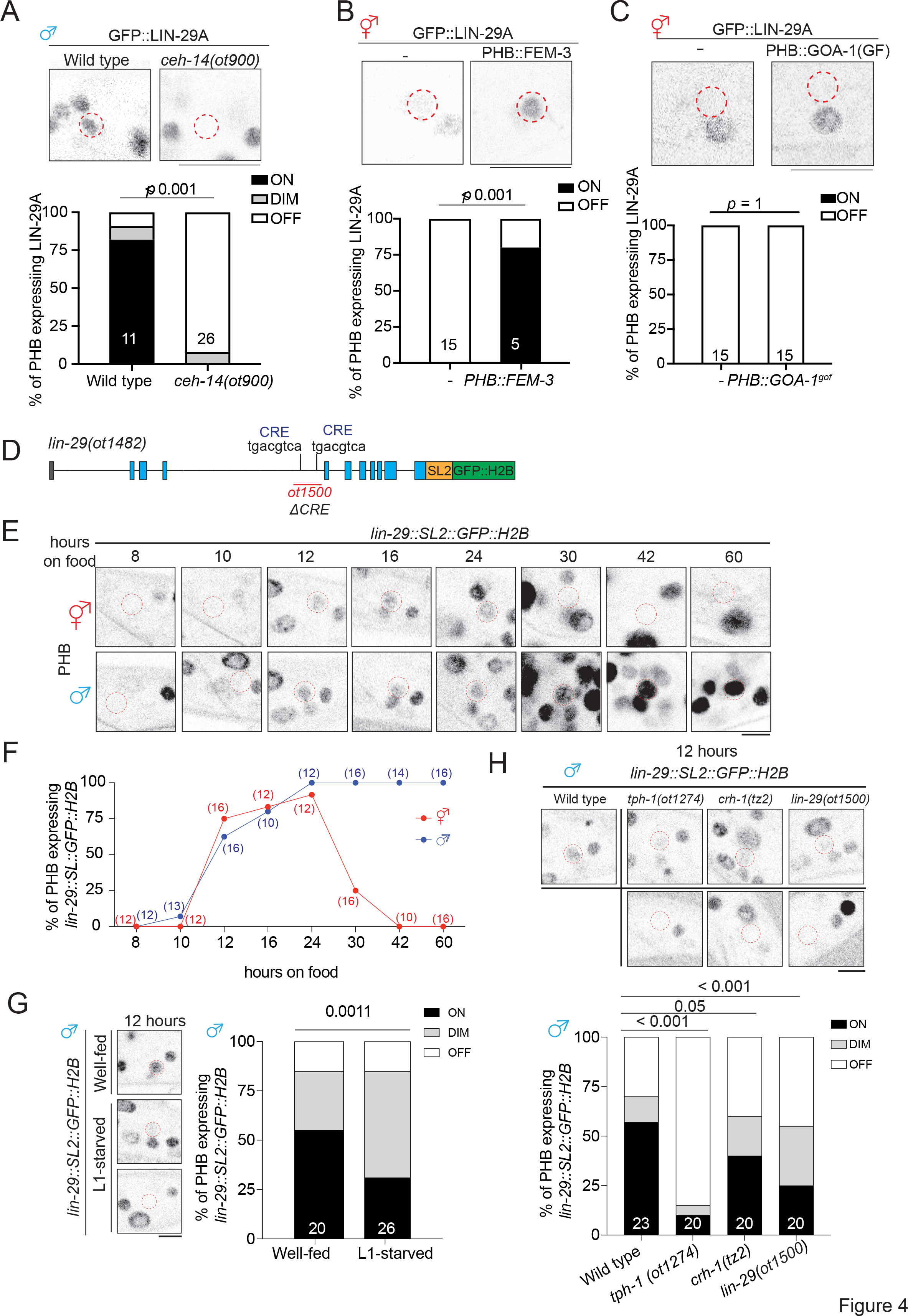
Early larval *lin-29a* transcription is regulated by serotonin>CREB signaling. **(A)** Representative images (top) and quantification of (bottom) PHB neurons expressing *lin- 29(xe63[gfp::lin-29a])* in wild type and *ceh-14(ot900)* males. **(B)** Representative images (top) and quantification of (bottom) PHB expressing *lin- 29(xe63[gfp::lin-29a])* with transgene masculinizing PHB (PHB::FEM-3)(*otEX7916*) in hermaphrodites. **(C)** Representative images (top) and quantification of (bottom) PHB expressing *lin- 29(xe63[gfp::lin-29a])* in hermaphrodites with transgene overexpressing GOA-1^gof^ (*otEx8037*) in the PHB. **(D)** Schematic illustration of *lin-29(ot1482[lin-29::SL2::GFP::H2B]) and lin-29(ot1500).* The *lin-29(ot1500)* allele is designed to delete the trunk of DNA elements, including the potential CRE site at the intron 3 of the *lin-29a* locus in *lin-29(ot1482*). **(E,F)** Longitudinal analysis of *lin-29(ot1482[lin-29::SL2::GFP::H2B])* expression in PHB neuron. Representative images (B) and quantification (C) of expression of PHB neuron expressing *lin-29(ot1482[lin-29::SL2::GFP::H2B])* in different time points after food exposure. **(G)** Representative images (left) and quantification of (right) PHB neurons expressing *lin- 29(ot1482[lin-29::SL2::GFP::H2B])* in males that undergo L1-starvation after 12 hours of food exposure. **(H)** Representative images (left) and quantification of (right) PHB neurons expressing *lin- 29(ot1482[lin-29::SL2::GFP::H2B])* in wild type, *tph-1(ot1274)*, *crh-1(tz2)*, and *lin-29(ot1500)* males after 12 hours of food exposure. Statistics: *chi*-squared tests followed by Bonferroni multiple comparisons test. *p*-value and N numbers are indicated on the graph. The red dashed circle indicates PHB. Scale bar = 5 µm. + indicates the mean value.

Previous work has shown that the temporal aspect of LIN-29A protein accumulation is controlled by the global heterochronic pathway, such that the translational inhibitor LIN-41 represses *lin-29a* translation in all cells until the fourth larval stage (AESCHIMANN *et al*. 2017; PEREIRA *et al*. 2019). We therefore surmised that the feeding state is bookmarked by CREB on the level of *lin-29a* transcription at earlier larval stages, to then set the stage for translational inhibition by the heterochronic pathway. To probe this issue, we measured *lin- 29a* gene transcription by generating an SL2-based transcriptional *lin-29a* reporter through CRISPR/Cas9 genome engineering (**Figure 4D**). We found that *lin-29a* transcription in PHB was induced in both sexes after animals were exposed to food for 12 hours (late L1 stage) and peaked at 24 hours (late L2 stage) (**Figure 4E,F, S3**). *lin-29a* transcription in another neuron that expresses LIN-29a protein in the adult, AVA, is not yet observed during these stages (**Figure S2B,C, S3**). In the hermaphrodite, *lin-29a* transcription began to be inhibited at the L3 stage, coinciding with the time when neuronal TRA-1 expression increased (BAYER *et al*. 2020b). The onset of *lin-29a* transcription after 12 hours of feeding in the late stage is reduced if animals have been starved prior to food exposure (**Figure 4G**). Similarly, animals lacking either *tph-1*, *crh-1*, or the CRE site in the *lin-29a* transcriptional reporter allele show reduced *lin-29a* transcription in PHB (**Figure 4H**). Taken together, LIN-29A in the PHB acts as a hub by integrating not only temporal (heterochronic pathway), sexual (TRA-1), and spatial (i.e. cell-type specific) information (CEH-14), but also an environmental axis that bookmarks past feeding status via CREB activation.

### LIN-29A is required to specify sexually dimorphic PHB>AVA synaptic connectivity

Having shown that early-life serotonin signaling regulates LIN-29A expression, we next asked if *lin-29a* mutant males phenocopied L1-starvation effects on sexually dimorphic synaptic connectivity. We found that the sexually dimorphic nature of PHB>AVA synapses was indeed abolished by two independent *lin-29a* null alleles, *xe38* and *xe40.* These defects can be measured with a GRASP transgene, as well as presynaptic CLA-1 and postsynaptic AVR-14 markers (**Figure 5A, B, C**; **Figure S4**). Sexually dimorphic connectivity of other LIN- 29-expressing neurons is not affected (**Figure S5**). Consistent with the L1 starvation results, PHB/AVA neurite contact length was not affected in *lin-29a* mutant animals (**Figure 5D,E**).

**Figure 5.**
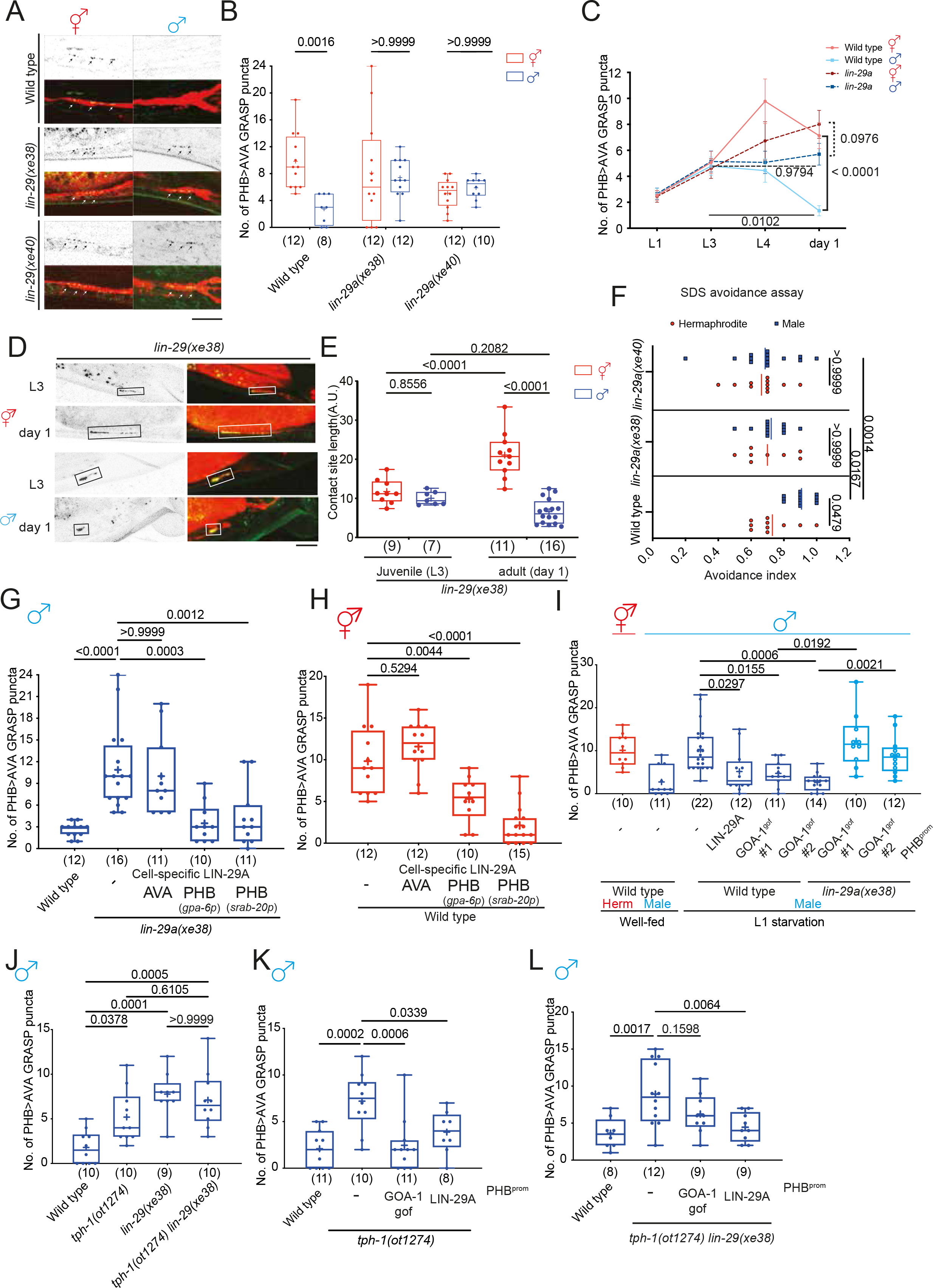
LIN-29A in PHB is required and sufficient to establish PHB>AVA sexual dimorphic connectivity upon sexual maturation. **(A,B)** Representative images (A) and quantification (B) of PHB>AVA synaptic GRASP(*otIs839*) in day 1 wild-type, *lin-29(xe38)* and *lin-29(xe40)* in both sexes. **(C)** Developmental analyses of PHB>AVA synaptic connectivity in wild-type and *lin-29a* mutants. N > 10 for each genotype and sex at any given time point. **(D)** Representative images of PHB>AVA neurite CD4-GRASP(*otEx8152*) in *lin-29a* mutants in both sexes. **(E)** Quantification of CD4-GRASP(*otEx8152*) *lin-29a* mutants in both sexes. **(F)** Quantification of SDS-avoidance assay in wild-type, *lin-29(xe38)* and *lin-29(xe40)*. **(G)** Quantification of PHB>AVA synaptic GRASP in *lin-29(xe38)* males with transgenes expressing LIN-29A cDNA in either AVA(*otEx7763*) or PHB (*otEx7790* for *gpa-6p* and *otEx7915* for *srab-20p*). **(H)** Quantification of PHB>AVA synaptic GRASP in wild-type hermaphrodite with transgenes expressing LIN-29A cDNA in either AVA(*otEx7763*) or PHB (*otEx7790* for *gpa-6p* and *otEx7915* for *srab-20p*). **(I)** Quantification of PHB>AVA synaptic GRASP(*otIs839*) in well-fed wild-type animals and L1-starved wild-type and *lin-29(xe38)* animals expressing PHB::LIN-29A (*otEx7915* ) or PHB::GOA-1^gof^(*otEx7925* and *otEx8158*). **(J)** Epistasis analysis of *tph-1* and *lin-29a* for PHB>AVA synaptic GRASP (*otIs839*) in males. **(K,L)** Quantification of PHB>AVA synaptic GRASP in *tph-1(ot1274)* (K) *and tph-1(ot1274) lin- 29(xe38)* (L) males with transgene overexpressing GOA-1^gof^ (*otEx7925)* and LIN- 29A(*otEx7915* ) in the PHB. Statistics: (B,E,F,I) Two-way ANOVA , (C) three-way ANOVA, and (G,H,J,K,L) one-way ANOVA followed by Bonferroni multiple comparisons test. *p*-value and N numbers are indicated on the graph. Scale bar = 10 µm. + indicates the mean value.

Since a *lin-29a* reporter allele is expressed in both presynaptic PHB and postsynaptic AVA (PEREIRA *et al*. 2019), we addressed the cellular focus of action of LIN-29A through cell- specific rescue experiments and found that restoration of LIN-29A in only the PHB neurons but not the AVA neurons restored proper synaptic elimination in *lin-29a* mutant males (**Figure 5G**). Moreover, overexpressing LIN-29A in the PHB neuron in wild-type hermaphrodites caused ectopic PHB>AVA synaptic loss (**Figure 5H**). We corroborated that *lin-29a* functions in PHB downstream of early juvenile food experience by showing that the rescuing effect of GOA-1^gof^ in starved males genetically depends on *lin-29a* (**Figure 5I**). Furthermore, genetic removal of *lin-29a* in serotonin-deficient *tph-1* mutant males did not further increase the synaptic defects compared to that of either *tph-1* or *lin-29a* single mutant animals (**Figure 5J**), Overexpression of either GOA-1^gof^ or LIN-29A in the PHB rescued the defects in serotonin-deficient *tph-1* males (**Figure 5K**), while overexpression of LIN-29A, but not GOA- 1^gof^ rescued the defects in *tph-1 lin-29a* double mutants (**Figure 5L**). Lastly, the effect of masculinization of PHB through PHB-specific TRA-1 removal on PHB>AVA connectivity (OREN-SUISSA *et al*. 2016) is suppressed by *lin-29a* removal (**Figure S6A**) but not the effect of AVA masculinization (**Figure S6B**). Taken together, our results suggest that LIN-29A in the PHB sensory neurons acts downstream of early juvenile food experience and is required and sufficient to promote synaptic elimination in males.

We investigated the structural requirements of the LIN-29A function. We found that the DNA binding activity of LIN-29A was required for its function since LIN-29A with Zn finger domain deletion failed to rescue the *lin-29a* defects and was insufficient to induce ectopic synaptic elimination (**Figure S7A,B**). We also found that the human homolog of LIN-29A, ZNF362, is able to rescue the synaptic elimination defects in *lin-29a* mutants (**Figure S7C**). Moreover, ZNF362 is also able to induce ectopic synaptic elimination in wild-type hermaphrodites, just as LIN-29A (**Figure S7D**).

We assessed the behavioral consequences of *lin-29a* by considering the physiological function PHB>AVA *en passant* synapses, which control *C. elegans* avoidance response to noxious chemicals such as SDS. This response is sexually dimorphic in day 1 adult animals such that hermaphrodites avoid SDS less than their male counterparts due to sex-specific PHB>AVA connectivity. We quantified the avoidance response in *lin-29a* mutant males and found that it was indeed feminized (**Figure 5F**).

### LIN-29A represses DMD-4 in PHB to promote sexually dimorphic PHB>AVA connectivity

Several sex-specific features are mediated by phylogenetically conserved Doublesex/Mab-3-related transcription factors (DMRTs). We had previously shown that the DMD-4 protein, one of several *C. elegans* Doublesex homologs, is initially expressed in juvenile PHA and PHB neurons in both sexes but becomes selectively degraded in male PHA and PHB neurons upon sexual maturation (BAYER *et al*. 2020a). The mutually exclusive expression pattern of LIN-29A (males) and DMD-4 (hermaphrodites) in PHB led us to investigate whether *lin-29a* may control DMD-4 degradation in male PHB. We find that DMD- 4::GFP protein fails to be degraded in PHB in a *lin-29a* mutant background (**Figure 6A, B**).

**Figure 6.**
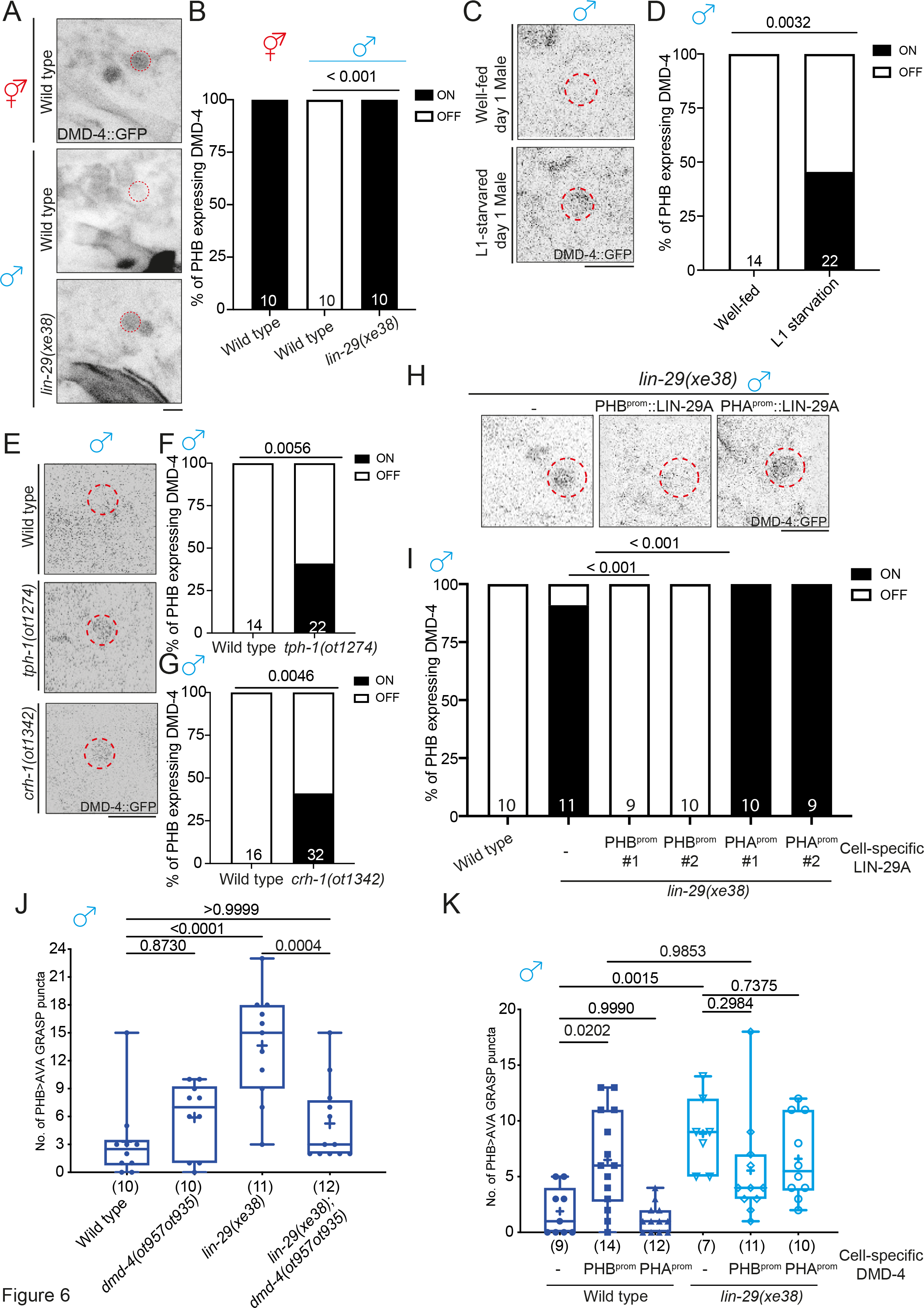
LIN-29A represses DMD-4 in PHB to control sexually dimorphic PHB>AVA connectivity. **(A,B)** Representative images (A) and (B) quantification of PHB neurons expressing *dmd- 4(ot935)* in wild-type hermaphrodite and male and *lin-29(xe38)* male. **(C,D)** Representative images of and (D) quantification of PHB neurons expressing *dmd- 4(ot935)* in well-fed and L1-starved day 1 males. **(E,F,G)** Representative images (E) and (F,G) quantification of PHB neurons expressing *dmd- 4(ot935)* in wild-type, *tph-1(ot1274)* and *crh-1(1342)* males. **(H)** (top) Representative images and (bottom) quantification of PHB neuron expressing *dmd- 4(ot935)* in *lin-29(xe38)* with transgenes that express LIN-29A cDNA in either PHB *(otEx7961 and otEx7964)* or PHA*(otEx8159 and otEx8160)*. **(J)** Epistasis mutant analysis of *lin-29a* and *dmd-4* for PHB>AVA synaptic GRASP *(otIs839)* in males. **(K)** Quantification of PHB>AVA synaptic GRASP *(otIs839)* in wild-type and *lin-29(xe38)* males with transgene overexpression DMD-4 in either PHB (*otEx7984*) or PHA(*otEx7983*). Statistics: (B,D,F,G,I) Two-proportion Z test, and (J,K) one-way ANOVA followed by Bonferroni multiple comparisons test. *p*-value and N numbers are indicated on the graph. The red dashed circle indicates PHB. Scale bar = 5 µm.

Since LIN-29A expression is feeding state-dependent, we predicted that L1 starvation (which leads to loss of LIN-29A expression) might stabilize DMD-4 protein expression in PHB and found this to be indeed the case (**Figure 6C,D**). Consistent with the implication of *tph-1* and *crh-1* in promoting *lin-29a* expression, DMD-4 expression in PHB was also stabilized in well- fed day 1 *tph-1* and *crh-1* mutant males (**Figure 6E-G**). Cell-specific rescue experiments showed that *lin-29a* functioned cell-autonomously in PHB to degrade DMD-4 in males (**Figures 6H, I**). We also find that PHB-specific expression of the human homolog of LIN-29A, ZNF362, rescues the effect of *lin-29a* on DMD-4 protein expression (**Figure S7E**).

As expected from DMD-4 expression in hermaphrodites (but not males), loss of *dmd-4* does not affect the lack of PHB>AVA synapse number growth in males (**Figure 6J**).

However, it is conceivable that it is the absence of *dmd-4* that accounts for PHB>AVA synapse number increase in males and that, hence, the derepression of DMD-4 in *lin-29a* mutants is responsible for the ectopic PHB>AVA synapses in males. To test this notion, we generated *lin-29a; dmd-4* double mutant animals and found that the synaptic defects found in *lin-29a* mutants were indeed suppressed (**Figure 6J**). Similarly, overexpressing DMD-4 in PHB neurons promoted the formation of *en passant* PHB>AVA synapses in males (**Figure 6K**). Loss of *lin-29a* did not further enhance the ectopic synaptic defect in males overexpressing DMD-4 in PHB, consistent with the epistatic relationship of these genes.

Taken together, *lin-29a* acts through DMD-4 to specify the sexually dimorphic nature of PHB>AVA *en passant* synapses.

### LIN-29A represses DMD-4 to inhibit *fmi-1* expression in adult male PHB

We identified a functionally relevant effector gene for the CREB>LIN-29>DMD-4 regulatory cassette through a nervous system-wide expression pattern analysis of putative synaptogenic molecules, including all members of the cadherin gene family (MM, CPL and OH, in prep.). We found that the unconventional cadherin protein FMI-1, the *C. elegans* homolog of the *Drosophila* Flamingo and vertebrate CELSR proteins (STEIMEL *et al*. 2010), showed a sexually dimorphic expression pattern in the PHB neurons of adult animals (**Figure 7A, B**). At juvenile stages, an SL2::GFP::H2B-based *fmi-1* reporter allele, generated by CRISPR/Cas9 genome engineering, showed non-dimorphic expression in PHB neurons, but upon sexual maturation, it was sex-specifically downregulated in males. Other neurons in vicinity to PHB show no sex-dependent difference (**Figure 7A**). Sexually dimorphic *fmi-1* expression is not only apparent at the transcriptional level (as measured with our SL2-based reporter allele) but can also be observed on the protein level. To visualize endogenous FMI-1 protein specifically in PHB, we engineered six copies of split GFP11(6xGFP11) at the C- terminus of *fmi-1* at the endogenous locus and overexpressed the other half of GFP with myristoylation peptide (myri-GPF1-10) in PHB (**Figure S8A,B)**. We found that reconstituted FMI-1::GFP intensity was significantly lower in day 1 males compared to their hermaphrodite counterparts, consistent with decreased *fmi-1* transcription in males upon sexual maturation (**Figure S8C,D**).

**Figure 7.**
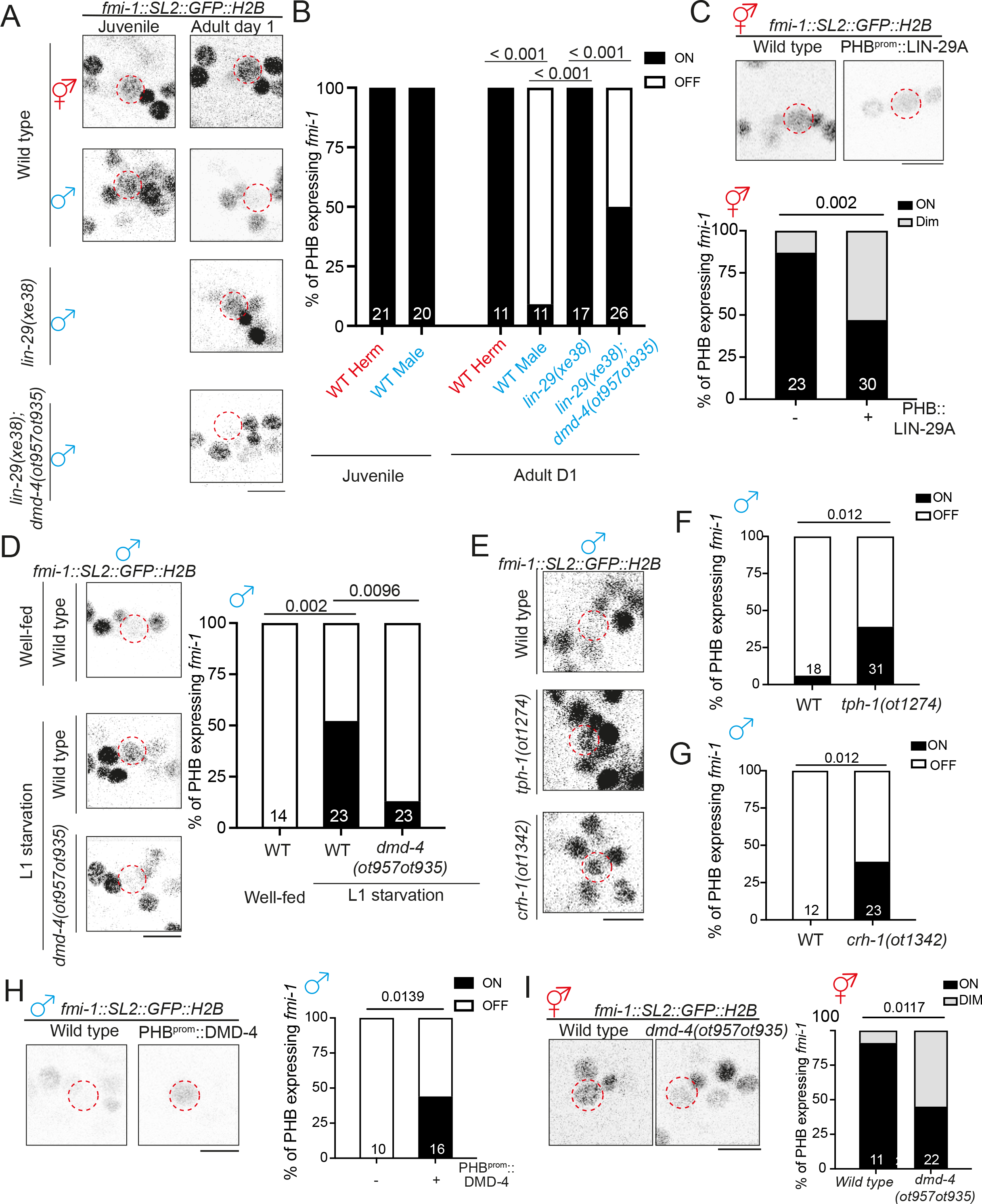
LIN-29A inhibits *fmi-1/Flamingo* expression via repressing DMD-4 **(A,B)** Representative images (A) and quantification (B) of *fmi-1(syb4563)* expression in the PHB in early L4 animals and day 1 animals with various genotypes. **(C)** Representative images (top) and quantification (bottom) of *fmi-1(syb4563)* expression in the PHB in wild-type hermaphrodites with transgene overexpressing LIN-29A cDNA(*otEx7961*) in the PHB neurons. **(D)** Representative images (left) and quantification (right) of *fmi-1(syb4563)* expression in the PHB in males that underwent L1 starvation. **(E,F,G)**Representative images (E) and quantification (F,G) of *fmi-1(syb4563)* expression in the PHB in wild-type, *tph-1(ot1274)* and *crh-1(ot1342)* males. **(H)** Representative images (top) and quantification (bottom) of *fmi-1(syb4563)* expression in the PHB in wild-type males with transgene overexpressing DMD-4 cDNA (*otEx8083*) in the PHB neurons. **(I)** Representative images (top) and quantification (bottom) of *fmi-1(syb4563)* expression in the PHB in *dmd-4(ot957ot935)* hermaphrodites. Statistics: (B,C,D,F,G,H,I) Two-proportion Z test, followed by Bonferroni multiple comparisons test. *p*-value and N numbers are indicated on the graph. The red dashed circle indicates PHB. Scale bar = 5 µm. + indicates the mean value.

Male-specific downregulation of *fmi-1* gene and FMI-1 protein expression was lost in *lin-29a* mutants (**Figure 7A,B; S8**). Expression of either LIN-29A in PHB or its vertebrate homolog, ZNF362, rescued the ectopic expression of FMI-1 in male PHB (**Figure S6F**). Ectopic expression of LIN-29A in hermaphrodite PHB shows that LIN-29A is not only required but also sufficient to downregulate *fmi-1* expression (**Figure 7C**). The feeding state- dependence of LIN-29A expression predicts that the downregulation of *fmi-1* should also be dependent on the feeding state. Indeed, we find that *fmi-1* expression in male PHB is derepressed upon either L1 starvation, or in well-fed serotonin-deficient *tph-1* or *crh-1/CREB* mutants (**Figure 7D-F**). The effect of *lin-29a* and feeding state on *fmi-1* expression is mediated by the *dmd-4* gene since the ectopic expression of *fmi-1* in male PHB in *lin-29a* mutants or after L1 starvation is suppressed by removal of *dmd-4* (**Figure 7D**). These findings indicate that *dmd-4* normally acts to promote *fmi-1* expression. Indeed, hermaphrodite-enriched PHB expression of *fmi-1* is reduced in *dmd-4* mutants and, conversely, overexpression of *dmd-4* in male PHB promotes *fmi-1* expression (**Figure 7H,I**).

### Separable functions of FMI-1 in controlling neurite contact length and synaptogenesis

The regulation of *fmi-1* by the serotonin>CREB>LIN-29A>DMD-4 axis, together with the documented role of vertebrate *fmi-1* orthologs in synaptogenesis (ZOU 2020), made us hypothesize that *fmi-1* may act as a synaptogenic molecule in the PHB>AVA context and that its serotonin/LIN-29A mediated downregulation suppresses PHB>AVA synapse number increases in males, hence generating synaptic sexual dimorphisms. Indeed, we found that a *fmi-1* null mutant allele that we generated by CRISPR/Cas9 genome engineering, results in decreased PHB>AVA GRASP puncta and presynaptic CLA-1 and postsynaptic AVR-14 puncta (**Figure 8A**; **Figure S9A-F**). Moreover, the PHB>AVA synaptic defects were rescued when FMI-1A was expressed in the PHB. In males, FMI-1A ectopic expression in the PHB also induced ectopic PHB>AVA synapses (**Figure 8B**). However, we also noted that the extent of PHB and AVA neurite contact, measured with CD4-based GRASP is significantly reduced as well (**Figure 8C**), already at the first larval stage (**Figure 8D**). The defect can be rescued by cell-specific re-expression in the PHB neurons (**Figure 8C,D**). This indicates that FMI-1 has, consistent with its role in other parts of the *C. elegans* nervous system (STEIMEL *et al*. 2010; NAJARRO *et al*. 2012), a role in neurite pathfinding and/or fasciculation of PHB during embryonic development, therefore preventing us from concluding that these synaptic defects are indeed the result of sexually dimorphic synaptogenic defects during postembryonic development.

**Figure 8.**
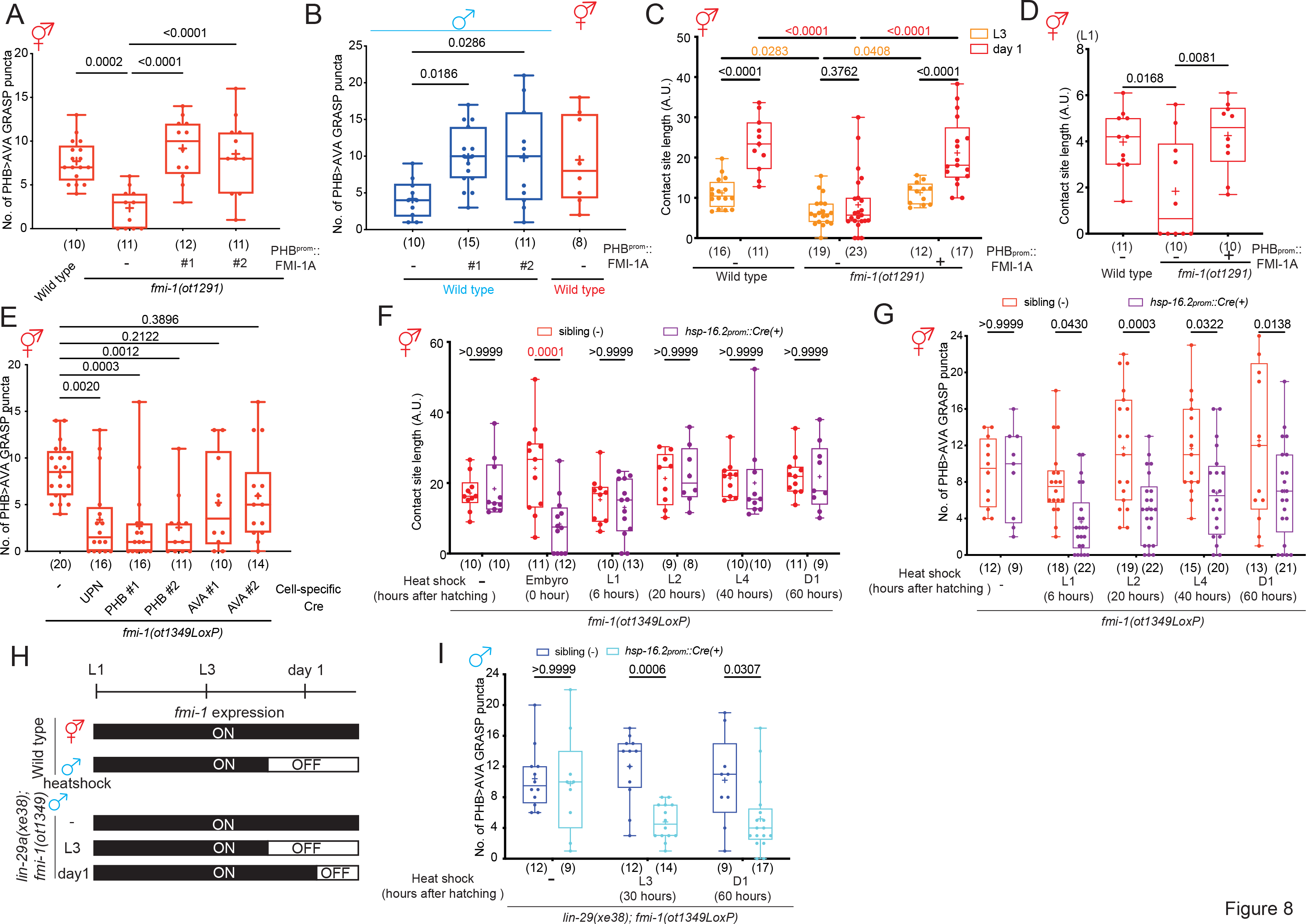
FMI-1 acts in PHB to promote the formation of *en passant* PHB>AVA synapses. **(A,B)** Quantification of PHB>AVA synaptic GRASP(*otIs839*) in *fmi-1(ot1291)* hermaphrodites (A) and wild-type males (B) with transgenes expressing FMI-1A cDNA (*otEx8032* and *otEx8033*)in the PHB neurons. **(C)** Quantification of CD4-GRASP(*otEx8152*) in L3 and day 1 wild-type, *fmi-1(ot1291)* and fmi-1(ot1291); PHB::FMI-1A. **(D)** Quantification of CD4-GRASP(*otEx8152*) in L1 wild-type, *fmi-1(ot1291)* and fmi- 1(ot1291); PHB::FMI-1A. **(E)** Quantification of PHB>AVA synaptic GRASP (*otIs839*) in *fmi-1(ot1349)* hermaphrodite with transgenes expressing Cre in pan-neuronally (*otEx8063*) or in PHB (*otEx8062* and *otEx8161*) or AVA(*otEx8064* and *otEx8162*). *fmi-1(ot1349)* is a *fmi-1* allele, which *fmi-1* locus is flanked with LoxP site and a GFP::H2B tagged at the C-terminus region. **(F)** Quantification of PHB>AVA synaptic GRASP (*otIs839*) in *fmi-1(ot1349)* hermaphrodites with transgenes expressing Cre in heat-shock promoter (*otEx8084*) and heat shock was performed by indicated time point. **(G)** Quantification of PHB>AVA synaptic GRASP (*otIs839*) in *fmi-1(ot1349)* hermaphrodites with transgenes expressing Cre in heat-shock promoter (*otEx8084*) and heat shock was performed by indicated time point. **(H)** Schematic illustration of *fmi-1* expression in the *lin-29(xe38); fmi-1(ot1349)* in heatshock *fmi-1* removal experiment. **(I)** Quantification of PHB>AVA synaptic GRASP (*otIs839*) in *lin-29(xe38);fmi-1(ot1349)* males with transgenes expressing Cre in heat-shock promoter (*otEx8084*) and heat shock was performed by indicated time point. Statistics: (A,B,D,E) One-way ANOVA and (C,F,G,I) two-way ANOVA followed by Bonferroni multiple comparisons test. *p*-value and N numbers are indicated on the graph. + indicates the mean value.

To separate embryonic from possible later roles of *fmi-1*, we employed a conditional gene removal strategy in which we inserted loxP sites into the reporter-tagged *fmi-1* locus (**Figure S8A**). We first confirmed that continuous removal of *fmi-1* using pan-neuronal and PHB-specific but not AVA-specific Cre driver lines recapitulated synaptic defects in *fmi- 1(ot1291)* null mutant (**Figure 8E**). We removed *fmi-1* in a temporally controlled removal manner using a heat-shock inducible Cre driver line. Embryonic induction of Cre expression recapitulated the neurite contact length defects, while *fmi-1* removal at any postembryonic stage had no effect on PHB/AVA contact length (**Figure 8F**). In contrast, eliminating *fmi-1* at postembryonic stages caused a decrease in PHB>AVA synapses in hermaphrodites compared to those of counterparts without transgene (**Figure 8G**). Moreover, we found that removal of *fmi-1* at the adult stage resulted in a reduction of PHB>AVA synapses, demonstrating that *fmi-1* is not only required for synapse number growth during sexual maturation, but is also continuously required to sustain synaptic connectivity in hermaphrodites.

Lastly, we asked whether ectopic *en passant* synapses in *lin-29a* mutant males result from derepressed synaptogenic *fmi-1* function in PHB. To this end, we removed *fmi-1* postembryonically in *lin-29a* mutant to bypass its critical embryonic neurite placement function. We found that removal of *fmi-1* in *lin-29a* mutant animals before sexual maturation (L3) and adulthood (day 1) significantly decreased the PHB>AVA synaptic puncta compared to the respective non-transgenic siblings **(Figure 8H and I)**.

Taken together these experiments demonstrate that FMI-1 has two separable functions, one during embryonic development in neurite outgrowth and placement and a synaptogenic one during postembryonic development. Since postembryonic expression of FMI-1 is sexually dimorphic, depending on the feeding state of the animal, we conclude that FMI-1 is the key synaptogenic effector gene of the CREB>LIN-29A>DMD-4 regulatory cascade. In hermaphrodites, this synaptogenic function is DMD-4-dependent and unimpeded by feeding state, while in male animals, feeding state and LIN-29A-dependent suppression of FMI-1 expression results in increase of sexually dimorphic *en passant* synapses.

## DISCUSSION

Early-life experience impacts later brain development or neurological disorders in a sex-specific manner, but the cellular and molecular mechanisms that lead to sex-specific vulnerability remain largely unknown. Here, we uncover the molecular basis of how the juvenile feeding-state experience affects the generation of sex-specificity of synaptic connectivity between a nociceptive sensory neuron and a command interneuron target.

A key regulatory bottleneck in this process is an evolutionarily conserved Zn finger transcription factor, LIN-29A, whose expression integrates four dimensions of specificity (**Fig.S10A**). Transcription of the *lin-29a* locus is directed to a subset of neuron cell types (like PHB) via terminal selectors, such as CEH-14 (shown here). The male-specificity of *lin-29a* transcription is imposed by the global sex identity regulator TRA-1, which antagonizes *lin-29a* transcription in hermaphrodites, while the temporal specificity of LIN-29A protein accumulation during sexual maturation is controlled by translational repression of *lin-29a* transcripts through the global heterochronic regulator *lin-41*. This repression is relieved upon sexual maturation by miRNA-mediated downregulation of *lin-41.* The fourth dimension of *lin- 29a* regulation is conferred by the feeding state of the animal. A serotonin- and CREB- dependent input is required specifically in the L1 stage to enable terminal selector-dependent induction of *lin-29a* transcription that is then translated into protein expression during sexual maturation via the relief of translational repression. Once the initial *lin-29a* transcription activation has been bookmarked, the locus becomes independent of a requirement for a feeding input (as evidenced by post-L1 starvation having no effect on LIN-29A expression).

Taken together, our studies tie an early transcriptional event, mediated by stimulus- dependent transcription factor, CREB, to the sustained expression of a locus, *lin-29a*, that later in life relays this input into the modulation of synaptic connectivity and, hence, information flow in the nervous system.

Our study here provides not only novel insights into the mechanistic basis of sculpting the sexually dimorphic nature of synaptic connectivity but reveals two distinct components of the establishment of sexually dimorphic connectivity: a sexually dimorphic increase in the adjacency of two neurites and a sexually dimorphic increase in the number of *en passant* synaptic connections. These two processes can be mechanistically uncoupled by by *fmi- 1/Flamingo*, which does not affect the postembryonic increase in neurite adjacency, but only affects the increase in *en passant* synapse number and, postdevelopmentally, maintains this synaptic connectivity. Considering the early function of *fmi-1* in controlling initial PHB/AVA neurite fasciculation during embryonic development, it is intriguing to note the lack of a role of *fmi-1* in controlling the sex-specific adjacency increase of the PHB and AVA neurites during postembryonic sexual maturation. This observation suggests that neurite contacts of the same two neurons can be regulated by distinct means during distinct stages of development. Moreover, the distinct functions of *fmi-1* in embryonic fasciculation and postembryonic synaptogenesis and maintenance are a likely reflection of context-dependent association of FMI-1 proteins with distinct interaction partners.

In vertebrates, the Flamingo orthologs CELSR1/2/3/4 have been implicated in axon outgrowth as well as synapse formation, but a function in maintaining synaptic structure had not been described before (TISSIR *et al*. 2005; LEWIS *et al*. 2011; FENG *et al*. 2012; CHAI *et al*. 2014; THAKAR *et al*. 2017; ZOU 2020). Our work uniquely places FMI-1 function, as well as its apparent dependence on past experience, in the context of sexually dimorphic synaptic connectivity. Vertebrate brains are thought to display sexually dimorphic features on multiple levels (SIMERLY 2002; MORRIS *et al*. 2004; DULAC AND KIMCHI 2007; YANG AND SHAH 2014; GEGENHUBER AND TOLLKUHN 2020), even though the cellular complexity of vertebrate brains has hampered the definition of such dimorphisms on a single neuron/single synapse level. We hope that our work will motivate a careful analysis of sexually dimorphic Flamingo expression in vertebrate brains, which may provide a critical entry point to not only identify vertebrate sexual dimorphisms but also understand their genetic specification.

## Supporting information

All Supp Figures

## ACKNOWLEDGEMENTS

We thank Chi Chen for generating transgenic strains and members of the Hobert lab, as well as Nathan Harris for comments on the manuscript. We thank Dr. Chun-Liang Pan for providing constructs for *fmi-1a* cDNA, *crh-1* variants and CD4-GRASP constructs and Dr. Meital Oren for the *avr-14* plasmid. Some of the strains were provided by the CGC, which is supported by the NIH (P40 OD010440). This work was supported by the NIH (R37NS039996) and the HHMI. Chien-Po Liao was supported by the Postdoctoral Research Abroad Program sponsored by the Ministry of Science and Technology from Taiwan and the Charles H. Revson Senior Fellowship in Biomedical Science (Grant No. 23-16).

## STAR methods

### KEY RESOURCE TABLE

| Bacterial and virus strains | Identifier | Resource |
| --- | --- | --- |
| *E. coli* | WormBase: OP50<br>WBStrain00041969 | CGC |
| **Chemicals, peptides, and recombinant proteins** |  |  |
| Alt-R S.p. Cas9 Nuclease V3 | Cat#1081059 | IDT |
| Alt-R CRISPR-Cas9 tracrRNA | Cat#1072533 | IDT |
| **Experimental models: Organisms/strains** |  |  |
| **Alleles** |  |  |
| *lin-29(xe38)* |  |  |
| *lin-29(xe40)* |  |  |
| *lin-29(xe63)* |  |  |
| *lin-29(ot1396)* |  |  |
| *lin-29(ot1482)* |  |  |
| *lin-29(ot1500)* |  |  |
| *tph-1(ot1274)* |  |  |
| *crh-1(tz2)* |  |  |
| *crh-1(ot1342)* |  |  |
| *him-8(e1489)* |  |  |
| *him-5(e1490)* |  |  |
| *fmi-1(ot1090)* |  |  |
| *fmi-1(ot1291)* |  |  |
| *fmi-1(ot1349)* |  |  |
| *fmi-1(ot1429)* |  |  |
| *fmi-1(syb4563)* |  |  |
| *dmd-4(ot935)* |  |  |
| *dmd-4(ot957ot935)* |  |  |
| **Strains** | **Identifier** | **Resource** |
| *lin-29(xe63); him-5(e1490); otEx8037[srab-20p::goa-1<sup>gof</sup>::SL2::TagRFP]* | OH18905 | This Study |
| *tph-1(ot1274) lin-29(xe63)/II; him-5(e1490)/V;otEx8037[srab-20p::goa-1<sup>gof</sup>::SL2::TagRFP]* | OH18389 | This Study |
| *tph-1(ot1274) lin-29(xe63)/II; crh-1(tz2); him-5(e1490)/V;otEx8037[srab-20p::goa-1<sup>gof</sup>::SL2::TagRFP]* | OH18390 | This Study |
| *lin-29(xe63)/II; crh-1(ot1342)/III;him-5(e1490)/V; otIs839* | OH18363 | This Study |
| *lin-29(xe63)/II; crh-1(tz2)/III; him-5(e1490)/V; otEx8037[srab-20p::goa-1<sup>gof</sup>::SL2::TagRFP]* | OH18413 | This Study |
| *tph-1(ot1274) lin-29(xe63)/II; him-5(e1490)/V; otEx8053[srab-20p::crh-1<sup>WT</sup>::SL2::TagRFP ]* | OH18439 | This Study |
| *tph-1(ot1274) lin-29(xe63)/II; him-5(e1490)/V;otEx8157[srab-20p::crh-1<sup>S48E</sup>::SL2::TagRFP ]* | OH18906 | This Study |
| *tph-1(ot1274) lin-29(xe63)/II; him-5(e1490)/V; otEx8113[srab-20p::crh-1<sup>S48A</sup>::SL2::TagRFP ]* | OH18636 | This Study |
| *lin-29(xe63)/II; crh-1(tz2)/III; him-5(e1490)/V; otEx8053[srab-20p::crh-1<sup>WT</sup>::SL2::TagRFP ]* | OH18907 | This Study |
| *lin-29(xe63)/II; crh-1(tz2)/III; him-5(e1490)/V; otEx8157[srab-20p::crh-1<sup>S48E</sup>::SL2::TagRFP ]* | OH18908 | This Study |
| *lin-29(xe63)/II; crh-1(tz2)/III; him-5(e1490)/V; otEx8113[srab-20p::crh-1<sup>S48A</sup>::SL2::TagRFP ]* | OH18909 | This Study |
| *lin-29(ot1396[ΔCRE])/II; him-5(e1490)/V; otEx8119[AVA::TagRFP, PHB::TagRFP]* | OH18669 | This Study |
| *lin-29(ot1482[lin-29::SL2::GFP::H2B])/II; him-5(e1490)/V;otIs839* | OH19115 | This Study |
| *tph-1(ot1274) lin-29(ot1482)/II; him-5(e1490)/V;otIs839* | OH19168 | This Study |
| *lin-29(ot1482)/II; crh-1(tz2)/III;him-5(e1490)/V;otIs839* | OH19169 | This Study |
| *lin-29(ot1500[ΔCRE]ot1482)him-5(e1490)/V;otIs839* | OH19177 | This Study |
| *him-5(e1490)/V;otIs839* | OH17170 | This Study |
| *lin-29(xe38)/II; him-5(e1490)/V;otIs839;* | OH18925 | This Study |
| *lin-29(xe40)/II; him-5(e1490)/V; otIs839;* | OH18926 | This Study |
| *otIs883; otEx8040[flp-18p(AVA)::TagRFP]* | OH18393 | This Study |
| *lin-29(xe38)/II; him-8(e1489)/IV; otIs883;*<br>*otEx8040[flp-18p(AVA)::TagRFP]* | OH18394 | This Study |
| *him-8(e1489)/IV; otIs902* | OH18703 | This Study |
| *lin-29(xe38)/II; him-8(e1489)/IV; otIs902* | OH18756 | This Study |
| *lin-29(xe38)/II; him-5(1490)/V; otIs839;*<br>*otEx7763[flp-18p::lin-29a::SL2::3xNLSGFP]* | OH17309 | This Study |
| *lin-29(xe38)/II; him-5(e1490)/V; otIs839;*<br>*otEx7915[srab-20p::lin-29a::SL2::3xNLS::GFP]* | OH17828 | This Study |
| *lin-29(xe38)/II; him-5(1490)/V; otIs839;*<br>*otEx7790[gpa-6p::lin-29a::SL2::3XNLS::GFP]* | OH18927 | This Study |
| *him-5(1490)/V; otIs839; otEx7790[gpa-6p::lin-29a::SL2::3XNLS::GFP]* | OH17411 | This Study |
| *him-5(1490)/V; otIs839; otEx7763[flp-18p::lin-29a::SL2::3xNLSGFP]* | OH18928 | This Study |
| *him-5(e1490)/V; otIs839; otEx7915[srab-20p::lin-29a::SL2::3xNLS::GFP]* | OH18929 | This Study |
| *him-5(e1490)/V; otIs839; otEx7925[srab-20p::goa-1<sup>gof</sup>::SL::3xNLS::GFP ]* | OH17867 | This Study |
| *him-5(e1490)/V; otIs839; otEx8158[srab-20p::goa-1<sup>gof</sup>::SL::3xNLS::GFP ]* | OH18911 | This Study |
| *lin-29(xe38)/II; him-5(e1490)/V; otIs839;*<br>*otEx7925[Psrab-20::goa-1(gof)::SL::3xNLS::GFP ]* | OH18930 | This Study |
| *lin-29(xe38)/II; him-5(e1490)/V; otIs839;*<br>*otEx8158[srab-20p::goa-1<sup>gof</sup>::SL::3xNLS::GFP ]* | OH18931 | This Study |
| *tph-1(ot1274); him-5(e1490)/V; otIs839* | OH18147 | This Study |
| *tph-1(ot1274) lin-29(xe38)/II; him-5(e1490)/V; otIs839* | OH18932 | This Study |
| *crh-1(tz2)/III ; him-5(e1490)/V; otIs839* | OH18217 | This Study |
| *crh-1(ot1342)/III; him-5(e1490)/V; otIs839* | OH18933 | This Study |
| *tph-1(ot1274); him-5(e1490)/V; otIs839;*<br>*otEx7925[srab-20p::goa-1<sup>gof</sup>::SL::3xNLS::GFP ]* | OH18934 | This Study |
| *tph-1(ot1274); him-5(e1490)/V; otIs839; otEx7915[srab-20p::lin-29a::SL2::3xNLS::GFP]* | OH18935 | This Study |
| *tph-1(ot1274) lin-29(xe38)/II; him-5(e1490)/V; otIs839; otEx7925[srab-20p::goa-1<sup>gof</sup>::SL::3xNLS::GFP]* | OH18936 | This Study |
| *tph-1(ot1274) lin-29(xe38)/II; him-5(e1490)/V; otIs839; otEx7915[srab-20p::lin-29a::SL2::3xNLS::GFP]* | OH18937 | This Study |
| *tph-1(ot1274); otIs839; him-5(e1490); otEx8045[srab-20p::crh-1<sup>WT</sup>::SL2::3xNLS::GFP]* | OH18415 | This Study |
| *tph-1(ot1274); otIs839; him-5(e1490); otEx8046[srab-20p::crh-1<sup>S48E</sup>::SL2::3xNLS::GFP]* | OH18416 | This Study |
| *tph-1(ot1274)/II; otIs839; him-5(e1490)/V; otEx8082[srab-20p::crh-1<sup>S48A</sup>::SL2::3xNLS::GFP]* | OH18504 | This Study |
| *crh-1(tz2) otIs839; him-5(e1490); otEx8045[srab-20p::crh-1<sup>WT</sup>::SL2::3xNLS::GFP]* | OH18912 | This Study |
| *crh-1(tz2) otIs839; him-5(e1490); otEx8046[srab-20p::crh-1<sup>S48E</sup>::SL2::3xNLS::GFP]* | OH18913 | This Study |
| *crh-1(tz2) otIs839; him-5(e1490)/V; otEx8082[srab-20p::crh-1<sup>S48A</sup>::SL2::3xNLS::GFP]* | OH18914 | This Study |
| *him-5(e1490)/V;dmd-4(ot935)/X; otIs839* | OH18938 | This Study |
| *lin-29(xe38)/II; him-5(e1490)/V;dmd-4(ot935)/X; otIs839* | OH18939 | This Study |
| *tph-1(ot1274)/II; him-5(e1490)/V;dmd-4(ot935)/X; otIs839* | OH18372 | This Study |
| *crh-1(ot1342)/III; him-5(e1490)/V;dmd-4(ot935)/X; otIs839* | OH18940 | This Study |
| *him-5(e1490) V; dmd-4(ot957ot935) X; otIs839* | OH18084 | This Study |
| *lin-29(xe38)/II; him-5(e1490) V; dmd-4(ot957ot935) X; otIs839* | OH18085 | This Study |
| *lin-29(xe38); him-5(e1490)/V; dmd-4(ot935)/X;otEx7961[srab-20p::lin-29a::SL2::TagRFP]* | OH18094 | This Study |
| *lin-29(xe38); him-5(e1490)/V; dmd-4(ot935)/X; otEx7964[srab-20p::lin-29a::SL2::TagRFP]* | OH18097 | This Study |
| *lin-29(xe38)/II; him-5(e1490)/V; dmd-4(ot935)/X; otEx7997[srab-20p::znf-362::SL2::TagRFP]* | OH18228 | This Study |
| *lin-29(xe38)/II; him-5(e1490)/V; dmd-4(ot935)/X; otEx7998[srab-20p::znf-362::SL2::TagRFP]* | OH18231 | This Study |
| *lin-29(xe38); him-5(e1490)/V; dmd-4(ot935)/X; otEx8159[srg-13p::lin-29a::SL2::TagRFP]* | OH18915 | This Study |
| *lin-29(xe38); him-5(e1490)/V; dmd-4(ot935)/X; otEx8160[srg-13p::lin-29a::SL2::TagRFP]* | OH18916 | This Study |
| *lin-29(xe38)/II; him-5(1490)/V; otIs839; otEx7984[gpa-6p::dmd-4::GFP]* | OH18186 | This Study |
| *lin-29(xe38)/II; him-5(1490)/V; otIs839; otEx7983[srg-13p::dmd-4::GFP]* | OH18185 | This Study |
| *him-5(1490)/V; otIs839; otEx7984[gpa-6p::dmd-4::GFP]* | OH18941 | This Study |
| *otIs839; otEx7983[srg-13p::dmd-4::GFP]* | OH18942 | This Study |
| *him-8(e1489)/IV; fmi-1(syb4563)/V; otIs839* | OH18943 | This Study |
| *lin-29(xe38)/II; him-8(e1489)/IV; fmi-1(syb4563)/V; otIs839* | OH18924 | This Study |
| *him-8(e1489); fmi-1(syb4563)/V; dmd-4(ot957ot935)/X; otIs839* | OH18272 | This Study |
| *lin-29(xe38); him-8(e1489); fmi-1(syb4563)/V; dmd-4(ot957ot935)/X; otIs839* | OH18982 | This Study |
| *him-8/IV; fmi-1(syb4563)/V; otEx7961[srab-20p::lin-29a::SL2::TagRFP]* | OH18944 | This Study |
| *tph-1(ot1274)/II; him-8(e1489)/IV; fmi-1(syb4563)/V; otIs839* | OH18945 | This Study |
| *crh-1(ot1342)/III; him-8(e1489)/IV; fmi-1(syb4563)/V; otIs839* | OH18946 | This Study |
| *lin-29(xe38)/II; him-8/IV; fmi-1(syb4563)/V; otEx7961[srab-20p::lin-29a::SL2::TagRFP]* | OH18985 | This Study |
| *him-8/IV; fmi-1(syb4563)/V; otEx8083[srab-20p::dmd-4::SL2::TagRFP]* | OH18505 | This Study |
| *him-8*/IV(e1489); *fmi-1*(ot1291)/V; *otIs839* | OH18271 | This Study |
| *lin-29*(xe38)/II; *him-8*(e1489)/IV; *fmi-1*(ot1291)/V;*otIs839* | OH18387 | This Study |
| *otIs839*;*him-5*(e1490); *otEx8032*[*srab-20p::fmi-1a::SL2::3xNLS::GFP*] | OH18370 | This Study |
| *otIs839*;*him-5*(e1490); *otEx8033*[*srab-20p::fmi-1a::SL2::3xNLS::GFP*] | OH18371 | This Study |
| *him-8*/IV(e1489); *fmi-1*(ot1291)/V; *otIs839*; *otEx8032*[*srab-20p::fmi-1a::SL2::3xNLS::GFP*] | OH19030 | This Study |
| *him-8*/IV(e1489); *fmi-1*(ot1291)/V; *otIs839*; *otEx8033*[*srab-20p::fmi-1a::SL2::3xNLS::GFP*] | OH19031 | This Study |
| *otIs839*; *him-5*(e1490) *fmi-1*(ot1349)/V | OH18441 | This Study |
| *otIs839*; *him-5*(e1490) *fmi-1*(ot1349)/V; *otEx8062*[*srab-20p::3xNLS::Cre*] | OH18461 | This Study |
| *otIs839*; *him-5*(e1490) *fmi-1*(ot1349)/V; *otEx8063*[*UPN::3xNLS::Cre*] | OH18467 | This Study |
| *otIs839*; *him-5*(1490)*fmi-1*(ot1349)/V;*otEx8064*[*flp-18p::3xNLS::Cre*] | OH18468 | This Study |
| *otIs839*; *him-5*(e1490) *fmi-1*(ot1349)/V;*otEx8084*[*hsp-16.2p::3xNLS::Cre*] | OH18506 | This Study |
| *otIs839*; *him-5*(e1490) *fmi-1*(ot1349)/V;*otEx8161*[*srab-20p::3xNLS::Cre*] | OH18917 | This Study |
| *otIs839*; *him-5*(1490)*fmi-1*(ot1349)/V;*otEx8162*[*flp-18p::3xNLS::Cre*] | OH18918 | This Study |
| *otIs839*; *him-5*(e1490) *fmi-1*(ot1349)/V;*otEx8084*[*hsp-16.2p::3xNLS::Cre*] | OH18506 | This Study |
| *him-8*(e1489)/IV;*otEx8152*[*srab-20p::TagRFP*, *srab-20p::CD4::GFP<sub>1-10</sub>*,*flp-18p::TagRFP*, *flp-18::CD4::GFP<sub>11</sub>*] | OH18858 | This Study |
| *him-8*(e1489)/IV; *fmi-1*(ot1291)/V;*otEx8152*[*srab-20p::TagRFP*, *srab-20p::CD4::GFP<sub>1-10</sub>*,*flp-18p::TagRFP*, *flp-18::CD4::GFP<sub>11</sub>*] | OH18860 | This Study |
| *him-8(e1489)/IV; fmi-1(ot1291)/V; otEx8152[srab-20p::TagRFP, srab-20p::CD4::GFP<sub>1-10</sub>, flp-18p::TagRFP, flp-18::CD4::GFP<sub>11</sub>], otEx8032[srab-20p::fmi-1a::SL2::3xNLS::GFP]* | OH18979 | This Study |
| *him-5(e1490) fmi-1(ot1349)/V; otEx8084[hsp-16.2p::3xNLS::Cre::p10UTR], otEx8152[srab-20p::TagRFP, srab-20p::CD4::GFP<sub>1-10</sub>, flp-18p::TagRFP, flp-18::CD4::GFP<sub>11</sub>]* | OH19023 | This Study |
| *lin-29(xe38)/II; otls839; him-5(e1490)/V; otEx7916[gpa-6p::fem-3::SL2::2xNLS::TagRFP-T]* | OH17830 | This Study |
| *otEx6829* | OH14590 | Cook et al., 2019. |
| *lin-29(xe38)/II; otls614; him-5(e1490)/V* | OH18553 | This Study |
| *otls614* | OH13577 | Oren-Suissa et al., 2016 |
| *otls630* | OH14099 | Oren-Suissa et al., 2016 |
| *him-8(e1489)/IV; otEx8176[srab-20p::gfp::cla-1 15ng/ul, flp-18p::avr-14::TagRFP]* | OH18983 | This Study |
| *him-8(e1489)/IV; fmi-1(ot1291)/V; otEx8176[srab-20p::gfp::cla-1 15ng/ul, flp-18p::avr-14::TagRFP]* | OH18984 | This Study |
| *lin-29(xe38)/II; him-8(e1489)/IV; otEx8176[srab-20p::gfp::cla-1 15ng/ul, flp-18p::avr-14::TagRFP]* | OH19008 | This Study |
| *lin-29(xe38)/II; otls839; him-5(e1490)/V; otEx7916[gpa-6p::fem-3::SL2::2xNLS::TagRFP-T]* | OH17830 | This Study |
| *otls839; him-5(e1490)/V; otEx8164[gpa-6p::fem-3::SL2::2xNLS::TagRFP-T]* | OH18920 | This Study |
| *otls839; him-5(e1490)/V; otEx8165[flp-18p::fem-3::SL2::2xNLS::TagRFP-T]* | OH18921 | This Study |
| *lin-29(xe38); otls839; him-5(e1490); otEx7929[srab-20p::lin-29a(delZn)::SL2::3xNLS::GFP]* | OH17984 | This Study |
| *otls839; him-5(e1490); otEx7928[srab-20p::lin-29a(delZn)::SL2::3xNLS::GFP]* | OH17881 | This Study |
| *lin-29(xe38);otl839; him-5(e1490); otEx7929[srab-20p::lin-29a(delZn)::SL2::3xNLS::GFP]* | OH17984 | This Study |
| *otls839; him-5(e1490)/V; otEx7930[srab-20p::znf362::SL2::3xNLS::GFP]* | OH17895 | This Study |
| *otls839; him-5(e1490)/V; otEx7931[srab-20p::znf362::SL2::3xNLS::GFP]* | OH17896 | This Study |
| *lin-29(xe38);otl839; him-5(e1490); otEx7929[srab-20p::lin-29a(delZn)::SL2::3xNLS::GFP]* | OH17984 | This Study |
| *lin-29(xe38)/II; him-5(e1490)/V; dmd-4(ot935)/X; otEx7997[srab-20p::znf-362::SL2::TagRFP]* | OH18228 | This Study |
| *lin-29(xe38)/II; him-5(e1490)/V; dmd-4(ot935)/X;otEx7998[srab-20p::znf-362::SL2::TagRFP]* | OH18231 | This Study |
| *tph-1(ot1274)/II;; him-8(e1489); otls902* | OH18769 | This Study |
| *him-5(e1490) fmi-1(ot1429)[fmi-1::6xGFP11]* | OH18838 | This Study |
| *him-5(e1490) fmi-1(ot1429)[fmi-1::6xGFP11];otEx8148[srab-20p::myriGFP::SL2::TagRFP]* | OH18839 | This Study |
| *lin-29(xe38)/II; him-5(e1490) fmi-1(ot1429)[fmi-1::6xGFP11];otEx8148[srab-20p::myriGFP::SL2::TagRFP]* | OH18887 | This Study |
| *otls839; him-5(e1490) fmi-1(ot1349)/V;otEx8084[Phsp-16.2::3xNLS::Cre]* | OH18506 | This Study |
| *lin-29(xe38)/II;fmi-1(ot1349) him-5(e1490)/V; otls839; otEx8084[hsp-16.2p::3xNLS::Cre]* | OH19231 | This Study |

### RESOURCE AVAILABILITY

#### Lead contact

For further detailed information and requests for resources and reagents, please contact Chien-Po Liao and Oliver Hobert

### EXPERIMENTAL DETAILS

#### Caenorhabditis elegans strains and handling

Worms were grown at 20°C on nematode growth media (NGM) plates seeded with E. coli (OP50) bacteria as a food source unless otherwise mentioned. Worms were maintained according to standard protocol. *him-8(e1489)* and *him-5(e1490)* were used as wild-type in this study to generate sufficient males. The key resources table lists a complete list of strains and transgenes generated and used in this study.

#### CRISPR/Cas9-based genome engineering

To generate *tph-1(ot1274)*, two guide crRNA (5’catcggatatctaaaagagg3’ and 5’ acctctcttcatctcaatat 3’) and ssODN (5’gtgccgaattccagaagcaccacgccatcggatatctaaaagaggccaacacaaagacacgttttcctgcagaagaggaa 3’) were used to remove the whole tph-1 locus. To generate *crh-1(ot1343)*, two guide crRNA (5’ taaggagattagttttccaa3’ and 5’ ttggagatttcttgttgagg 3’) and ssODN (5’ gtgtttgtttttcaaagaatagcttatatatatgatgaaatctcgtttttatttttatttcctaattttt 3’) were used to remove the whole *crh-1* locus. To generate lin-29(ot1396), two guide crRNA (5’ atttgaacccaatattgaat3’ and 5’ gagttcattttgatttcacg 3’) and ssODN (5’ agtggtcaaagaaatttgagagaaaaagtgcggagcgtgaaatcaaaatgaactcggctatatttcggcc 3’) were used to remove the potential CRE site in the *lin-29a* locus. To generate *fmi-1(ot1090)* and *fmi- 1(ot1291)*, two guide crRNA (5’ ttgaatgtgaatgtcagtgg3’ and 5’ TGATGCGTATTACACATATA 3’) were used to remove the whole *fmi-1* locus. To generate fmi-1(ot1349), crRNA (5’aagaactgaccagctgccaa3’) and ssODN (5’ ctgatacaaccctttgctcttttcacctcatatgtATAACTTCGTATAGCATACATTATACGAAGTTATcccgggttggcagctggtcagttcttcttccaaagagacgc3’) were used to insert the first LoxP site at 5’UTR region and second crRNA(5’ atcggaacaatgaacaagta 3’) and homemade LoxP::GFP::H2B ssODN via PCR and exonucleuase were used to insert the second LoxP site. To generate *fmi-1(ot1429)*, crRNA (5’ TACCACATCTACATTCAACA3’) and codon-optimized 6XGFP11 ssODN were used to remove the original stop codon and insert GFP11 at the C terminus of fmi-1 gene locus. To generate, lin-29(ot1482), crRNA (5’ ttatcggaatatgtgagttc3’) and homemade SL2::GFP::H2B ssOND were used.

#### Molecular cloning

To generate pCPL1(*srab-20p::goa-1^gof^::SL2::TagRFP*) and pCPL4(*srab-20p::goa- 1^gof^::SL2::3XNLS::GFP*) , upstream ∼1.2kb promoter from *srab-20*, which was amplified from (*srab-20p::GFP),* the synthetic *goa-1^gof^* DNA fragment, and the SL2-backbone amplified from *pEAB42 (srg-13p::ser-4::SL2::tagRFP)* were ligated. To generate pCPL2(*gpa-6p::lin- 29a::SL2::3XNLS::GFP*) and pCPL3(*srab-20p::lin-29a::SL2::3XNLS::GFP*), *srab-20p* and *gpa-6p* amplified from pCPL4 and pEAB3 (2.6 kb of the *gpa-6* promoter fused to GFP) respectively, and lin-29a cDNA (1.4kb) and the SL2-base backbone amplified from pCPL4 were ligated. To generate pCPL5 (*srab-20p::crh-1^WT^::SL2::3XNLS::GFP),* pCPL6 (*srab- 20p::crh-1^S48E^::SL2::3XNLS::GFP) and* pCPL7 (*srab-20p::crh-1^S48A^::SL2::3XNLS::GFP),* crh-1 variants subcloned from gcy-8*p::crh-1^WT^,* gcy-8*p::crh-1^S48E^, and* gcy-8*p::crh-1^S48A^* (gifts from Dr.Chun-Liang Pan) and backbone amplified from pCPL4 were ligated. To generate pCPL8 (*srab-20p::crh-1^WT^::SL2::TagRFP),* pCPL9 (*srab-20p::crh-1^S48E^::SL2::TagRFP) and* pCPL10 (*srab-20p::crh-1^S48A^::SL2::TagRFP), crh-1* variants subcloned into pCPL1 backbone. To generate pCPL11(*srab-20p::lin-29a::SL2::TagRFP*) and pCPL12(*srg-13p::lin- 29a::SL2::TagRFP*), lin-29a was subcloned into the backbone amplified from pCPL1 and pEAB42 (*srg-13p::ser-4::SL2::TagRFP*) respectively. To generate pCPL13(*srab-20p::dmd- 4::SL2::TagRFP*), *dmd-4* cDNA (0.8 kb) was amplified and subcloned into pCPL1. To generate pCPL14(*srab-20p::fmi-1a::SL2::3XNLS::GFP*), *fmi-1a* cDNA was amplified from *ser2prom3::fmi-1a* (gift from Dr. Chun-Liang Pan) and subcloned into pCPL4. To generate pCPL15 (*UPN::3xNLS::Cre*), pCPL16(*srab-20p::3xNLS::Cre*) pCPL17 (*flp-18p::3XNLS::Cre*) and pCPL23(*hsp16.2::3xNLS::Cre*), UPN, *srab-20p*, *flp-18p* (∼1.4kb) and *hsp16.2p* (∼0.4 kb) were subcloned into pCC301(*rab-3p1::3xNLS::Cre*). To generate pCPL18 (*srab-20p::gfp::cla- 1(s)*), *srab-20p* were digested from pCPL1 by SphI and XmaI restriction enzymes and ligated into the backbone of pMM13(*cat-4p::gfp::cla-1(s)*). To generate pCPL19 (*flp-18p::avr- 14::TagRFP*), *avr-14* was amplified from *avr-14* plasmid (gift from Dr. Meital Oren) and ligated into pCC45(*flp-18p::TagRFP*). To generate pCPL20(*srab-20p::znf362::SL2::3XNLS::GFP*) and pCPL21(*srab-20p::znf362::SL2::TagRFP*), synthetic codon-optimized human znf362 cDNA was subcloned into pCPL4 and pCPL1 respectively. To generate pCPL24(*srab- 20p::myriGFP1-10::SL2::TagRFP*), split GFP1-10 was cloned with myristylation sequences and ligated with pCPL1 backbone. To generate pCPL25 (*srab-20p::CD4::GFP1-10*) and pCPL26(*flp-18p::CD4::GFP11*), *srab-20p* and *flp-18p* were digested from pCPL16 and pCPL17 respectively, and ligated to the backbone from *ser2prom3p::CD4:: GFP1-10* and *unc- 17p::CD4:: GFP11* (gifts from Dr. Chun-Liang Pan), respectively.

#### L1 starvation assay

Animals with indicated genotype were synchronized by hypochlorite treatment of gravid adults followed by 12 hours in M9 at 20℃ for embryos to hatch. Synchronized L1 animals were released onto the unseeded NGM plates for another 24 hours. L1-starved animals were then washed and transferred to seeded NGM plates.

#### Heat shock assay

OH18506, *him-5(e1490) fmi-1(ot1349); otIs839, otEx8084[hsp-16.2p::3xNLS::Cre::p10UTR],* OH19023*, him-5(e1490) fmi-1(ot1349)/V;otEx8084[hsp- 16.2p::3xNLS::Cre::p10UTR],otEx8152[srab-20p::TagRFP, srab-20p::CD4::GFP1-10,flp-18p::TagRFP, flp-18::CD4::GFP11],* OH19231, *lin-29(xe38);fmi-1(ot1349) him-5(e1490); otIs839; otEx8084[hsp-16.2p::3xNLS::Cre]*,animals were synchronized by hypochlorite treatment of gravid adults followed by 12 h ours in M9 at 20℃ for embryos to hatch.

Synchronized L1 animals were released onto the seeded NGM plates, and heat shock was performed 6, 20, 40 and 60 hours after releasing onto the seeded plates (indicates as hours after hatching the figures). Animals were heat shocked at 34°C for 20 minutes, followed by 20 minutes of resting at 20°C three times to induce sufficient heat shock response. PHB>AVA GRASP puncta and PHB/AVA adjacency were analyzed at day 1 stage for groups that received heat shock at 6, 20, or 40 hours post-hatching. For groups that received heat shock at 60 hours post-hatching, analyses of PHB>AVA GRASP puncta and PHB/AVA adjacency were conducted 24 hours after heat shock.

#### SDS avoidance behavior

The SDS avoidance assay was based on procedures as described. To deliver the testing droplets, we pulled 10-μl glass capillary pipette (VWR international) by hand on the flame to reduce the diameter of the tip and mounted the capillary pipette on a rubber tubing and operated by mouth. We delivered a small drop of solution containing either the repellent (0.1% SDS in M13 buffer) or buffer (M13 buffer: 30 mM Tris-HCl pH 7.0, 100 mM NaCl, 10mM KCl) to near the tail of an animal while it moves forward. Once in contact with the tail, the drop surrounded the animal by capillary action and reached the anterior head region.

Assayed worms were transferred individually to fresh and unseeded NGM plates. Each assay started by testing the animals with drops of M13 buffer alone. The response to each drop was scored as reversing or not reversing. The avoidance index is the number of reversal responses divided by the total number of trials. An interstimulus interval of at least two minutes was used between successive drops of the same animal.

#### Microscopy

Worms were anesthetized in 100 mM of sodium azide on the 5% agarose on glass slides. All images were acquired using a Zeiss confocal microscope (LSM 880 or LSM980).For synaptic GRASP and gene expression experiments, animals were imaged using 63 X objective and with a fixed imaging setting. For CLA-1 and AVR-14 puncta experiments, animals were imaged using 40 X objective and with a fixed imaging setting. For PHB-AVA adjacency CD4- GRASP experiments, animals were 40 X objective and with a fixed imaging setting.

### QUANTIFICATION AND STATISTICAL ANALYSIS

#### Quantification of synaptic GRASP puncta

For all the synaptic GRASP, the images were acquired using 63 X objective and with a fixed imaging setting either with LSM880 or LSM980. The raw images were unbiasedly analyzed with PysQi (MAJEED *et al*. 2024), the automatic puncta quantification software.

#### Quantification of synaptic iBLINC puncta

For iBLINC experiments, animals were imaged using a 63 X objective with a fixed imaging setting with LSM880, and puncta were quantified by scanning the original full Z-stack for distinct dots in the area where the processes of the two neurons overlap.

#### Quantification of GFP::LIN-29A, DMD-4::GFP, and *lin-29a* and *fmi-1* expression

For expression of translational and transcriptional *lin-29a* reporter constructs, images of *lin- 29(xe63[gfp::lin-29a])* and *lin-29(ot1482[lin-29::SL2::GFP::H2B])*, animals with different mutant or transgene overexpression backgrounds were acquired using 63 X objective with fixed imaging settings with either LSM880 or LSM980. The expression level is categorized into three tiers: on, dim, and off. Cells with GFP fluorescent intensity lower than 50% of the normal “on” cells are identified as “dim.”

For DMD-4::GFP quantification, images of *dmd-4(ot935)* animals with different mutant or transgene overexpression backgrounds were acquired using 63 X objective with fixed imaging settings with either LSM880 or LSM980.

For fmi-1 gene expression quantification, images of *fmi-1(syb4563)* animals with different mutant or transgene overexpression backgrounds were acquired using 63 X objective with fixed imaging settings with either LSM880 or LSM980. Cells with GFP fluorescent intensity lower than 50% of the normal “on” cells are identified as “dim.”

#### Quantification of PHB and AVA contact CD4 GRASP

The contact site length between PHB and AVA processes in the CD4 reporter was quantified in Fiji ImageJ (SCHINDELIN *et al*. 2012). Briefly, the entire Z-stack was scanned while tracing over the GFP+ region (where the PHB and AVA processes overlap) with a segmented line and then measuring the overall line length. In cases where the contact and resulting GFP signal was discontinuous, multiple lines were drawn, measured independently, and summed to yield the overall contact site length. For visualization purposes, figures contain a representative subset of the Z-stack reconstructed as maximum intensity projection using Zeiss Zen software to display the maximal PHB-AVA contact site.

#### Quantification of CLA-1 and AVR-14 puncta

GFP::CLA-1 and AVR-14::TagRFP puncta in the PHB and AVA, respectively, were quantified manually in Fiji by scanning the entire Z-stack and only scoring puncta co-localizing with cytoplasmic AVAp::RFP and cytoplasmic PHBp::GFP, respectively.

#### Quantification of the juxtaposition of the CLA-1 and AVR-14 puncta

For scoring the juxtaposition of the PHB GFP::CLA-1 and AVA AVR-14::RFP puncta, each Z- stack was first scanned in the region of interest to quantify all GFP::CLA-1 puncta. Next, AVR-14::TagRFP puncta directly adjacent to with CLA-1 puncta were scored. The juxtaposition index is calculated as follows: AVR-14::TagRFP juxtaposed with CLA-1::GFP/Total GFP::CLA-1)*100%.

## REFERENCES

1. Aeschimann, F., P. Kumari, H. Bartake, D. Gaidatzis, L. Xu et al., 2017 LIN41 Post- transcriptionally Silences mRNAs by Two Distinct and Position-Dependent Mechanisms. Mol Cell 65: 476–489 e474.

2. Bayer, E. A., and O. Hobert, 2018 Past experience shapes sexually dimorphic neuronal wiring through monoaminergic signalling. Nature 561: 117–121.

3. Bayer, E. A., R. C. Stecky, L. Neal, P. S. Katsamba, G. Ahlsen et al., 2020a Ubiquitin- dependent regulation of a conserved DMRT protein controls sexually dimorphic synaptic connectivity and behavior. Elife 9.

4. Bayer, E. A., H. Sun, I. Rafi and O. Hobert, 2020b Temporal, Spatial, Sexual and Environmental Regulation of the Master Regulator of Sexual Differentiation in C. elegans. Curr Biol 30: 3604–3616 e3603.

5. Bockaert, J., S. Claeysen, C. Becamel, A. Dumuis and P. Marin, 2006 Neuronal 5-HT metabotropic receptors: fine-tuning of their structure, signaling, and roles in synaptic modulation. Cell Tissue Res 326: 553–572.

6. Chai, G., L. Zhou, M. Manto, F. Helmbacher, F. Clotman et al., 2014 Celsr3 is required in motor neurons to steer their axons in the hindlimb. Nat Neurosci 17: 1171–1179.

7. Cook, S. J., T. A. Jarrell, C. A. Brittin, Y. Wang, A. E. Bloniarz et al., 2019 Whole-animal connectomes of both Caenorhabditis elegans sexes. Nature 571: 63–71.

8. Cook, S. J., C. A. Kalinski and O. Hobert, 2023 Neuronal contact predicts connectivity in the C. elegans brain. Curr Biol 33: 2315–2320 e2312.

9. Dulac, C., and T. Kimchi, 2007 Neural mechanisms underlying sex-specific behaviors in vertebrates. Curr Opin Neurobiol 17: 675–683.

10. Feinberg, E. H., M. K. Vanhoven, A. Bendesky, G. Wang, R. D. Fetter et al., 2008 GFP Reconstitution Across Synaptic Partners (GRASP) defines cell contacts and synapses in living nervous systems. Neuron 57: 353–363.

11. Feng, J., Y. Xu, M. Wang, Y. Ruan, K. F. So et al., 2012 A role for atypical cadherin Celsr3 in hippocampal maturation and connectivity. J Neurosci 32: 13729–13743.

12. Gat, A., V. Pechuk, S. Peedikayil-Kurien, S. Karimi, G. Goldman et al., 2023 Integration of spatially opposing cues by a single interneuron guides decision-making in C. elegans. Cell Rep 42: 113075.

13. Gegenhuber, B., and J. Tollkuhn, 2020 Signatures of sex: Sex differences in gene expression in the vertebrate brain. Wiley Interdiscip Rev Dev Biol 9: e348.

14. Gurel, G., M. A. Gustafson, J. S. Pepper, H. R. Horvitz and M. R. Koelle, 2012 Receptors and other signaling proteins required for serotonin control of locomotion in Caenorhabditis elegans. Genetics 192: 1359–1371.

15. Herringa, R. J., R. M. Birn, P. L. Ruttle, C. A. Burghy, D. E. Stodola et al., 2013 Childhood maltreatment is associated with altered fear circuitry and increased internalizing symptoms by late adolescence. Proc Natl Acad Sci U S A 110: 19119–19124.

16. Hiscox, L. V., T. H. Sharp, M. Olff, S. Seedat and S. L. Halligan, 2023 Sex-Based Contributors to and Consequences of Post-traumatic Stress Disorder. Curr Psychiatry Rep 25: 233–245.

17. Honeycutt, J. A., C. Demaestri, S. Peterzell, M. M. Silveri, X. Cai et al., 2020 Altered corticolimbic connectivity reveals sex-specific adolescent outcomes in a rat model of early life adversity. Elife 9.

18. Jarrell, T. A., Y. Wang, A. E. Bloniarz, C. A. Brittin, M. Xu et al., 2012 The connectome of a decision-making neural network. Science 337: 437–444.

19. Kagoshima, H., G. Cassata, Y. G. Tong, N. Pujol, G. Niklaus et al., 2013 The LIM homeobox gene ceh-14 is required for phasmid function and neurite outgrowth. Dev Biol 380: 314–323.

20. Kimura, Y., E. E. Corcoran, K. Eto, K. Gengyo-Ando, M. A. Muramatsu et al., 2002 A CaMK cascade activates CRE-mediated transcription in neurons of Caenorhabditis elegans. EMBO Rep 3: 962–966.

21. Lewis, A., N. Wilson, G. Stearns, N. Johnson, R. Nelson et al., 2011 Celsr3 is required for normal development of GABA circuits in the inner retina. PLoS Genet 7: e1002239.

22. Li, Z., J. Zhou, K. A. Wani, T. Yu, E. A. Ronan et al., 2023 A C. elegans neuron both promotes and suppresses motor behavior to fine tune motor output. Front Mol Neurosci 16: 1228980.

23. Lonze, B. E., and D. D. Ginty, 2002 Function and regulation of CREB family transcription factors in the nervous system. Neuron 35: 605–623.

24. Lupien, S. J., B. S. McEwen, M. R. Gunnar and C. Heim, 2009 Effects of stress throughout the lifespan on the brain, behaviour and cognition. Nat Rev Neurosci 10: 434–445.

25. Majeed, M., H. Han, K. Zhang, W. X. Cao, C. P. Liao et al., 2024 Toolkits for detailed and high-throughput interrogation of synapses in C. elegans. Elife 12.

26. Morris, J. A., C. L. Jordan and S. M. Breedlove, 2004 Sexual differentiation of the vertebrate nervous system. Nat Neurosci 7: 1034–1039.

27. Najarro, E. H., L. Wong, M. Zhen, E. P. Carpio, A. Goncharov et al., 2012 Caenorhabditis elegans Flamingo Cadherin fmi-1 Regulates GABAergic Neuronal Development. J Neurosci 32: 4196–4211.

28. Oren-Suissa, M., E. A. Bayer and O. Hobert, 2016 Sex-specific pruning of neuronal synapses in Caenorhabditis elegans. Nature 533: 206–211.

29. Oury, F., V. K. Yadav, Y. Wang, B. Zhou, X. S. Liu et al., 2010 CREB mediates brain serotonin regulation of bone mass through its expression in ventromedial hypothalamic neurons. Genes Dev 24: 2330–2342.

30. Pereira, L., F. Aeschimann, C. Wang, H. Lawson, E. Serrano-Saiz et al., 2019 Timing mechanism of sexually dimorphic nervous system differentiation. Elife 8.

31. Schindelin, J., I. Arganda-Carreras, E. Frise, V. Kaynig, M. Longair et al., 2012 Fiji: an open- source platform for biological-image analysis. Nat Methods 9: 676–682.

32. Serrano-Saiz, E., Richard J. Poole, T. Felton, F. Zhang, Estanisla D. De La Cruz et al., 2013 Modular Control of Glutamatergic Neuronal Identity in C. elegans by Distinct Homeodomain Proteins. Cell 155: 659–673.

33. Simerly, R. B., 2002 Wired for reproduction: organization and development of sexually dimorphic circuits in the mammalian forebrain. Annu Rev Neurosci 25: 507–536.

34. Steimel, A., L. Wong, E. H. Najarro, B. D. Ackley, G. Garriga et al., 2010 The Flamingo ortholog FMI-1 controls pioneer-dependent navigation of follower axons in C. elegans. Development 137: 3663–3673.

35. Thakar, S., L. Wang, T. Yu, M. Ye, K. Onishi et al., 2017 Evidence for opposing roles of Celsr3 and Vangl2 in glutamatergic synapse formation. Proc Natl Acad Sci U S A 114: E610–E618.

36. Tissir, F., I. Bar, Y. Jossin, O. De Backer and A. M. Goffinet, 2005 Protocadherin Celsr3 is crucial in axonal tract development. Nat Neurosci 8: 451–457.

37. White, J. G., E. Southgate, J. N. Thomson and S. Brenner, 1986 The structure of the nervous system of the nematode *Caenorhabditis elegans*. Philosophical Transactions of the Royal Society of London B. Biological Sciences 314: 1–340.

38. Yang, C. F., and N. M. Shah, 2014 Representing sex in the brain, one module at a time. Neuron 82: 261–278.

39. Zhang, J., C. Y. Cai, H. Y. Wu, L. J. Zhu, C. X. Luo et al., 2016 CREB-mediated synaptogenesis and neurogenesis is crucial for the role of 5-HT1a receptors in modulating anxiety behaviors. Sci Rep 6: 29551.

40. Zou, Y., 2020 Breaking symmetry - cell polarity signaling pathways in growth cone guidance and synapse formation. Curr Opin Neurobiol 63: 77–86.

