## Supplementary material for "Experience-dependent, sexually dimorphic synaptic connectivity defined by sex-specific cadherin expression": All Supp Figures

### SUPPLEMENTARY FIGURES & LEGENDS

#### SUPPLEMENTARY FIGURE LEGENDS

**Figure S1. Serotonin signaling affects PHB>AVA synaptic connectivity. Related to Figure 1 and 2.**

**(A)** (top) Schematic illustration of the juxtaposition of GFP::CLA-1 in the PHB neuron and AVR-14::TagRFP in the AVA neuron in L3 and day 1 well-fed and L1-starved animals in both sexes. The juxtaposition index is calculated by the ratio of GFP::CLA-1 that juxtaposed to AVR-14::TagRFP to the total GFP::CLA-1 in the PHB. (bottom)Representative images of the juxtaposition of PHB-localized GFP::CLA-1 and AVA-localized AVR-14::TagRFP (*otEx8163*) in L3 and day 1 well-fed and L1-starved animals in both sexes.

**(B)** Quantification of juxtaposition index in L3 and day 1 well-fed and L1-starved animals in both sexes.

**(C)** Representative images (top) and quantification (bottom) of AVA-juxtaposed GFP::CLA-1 in PHB (*otIs883;otEx8040*) in day 1 wild-type and *tph-1(ot1274)*.

**(D)** Representative images (top) and quantification (bottom) of PHB-juxtaposed AVR-14::TagRFP (*otIs902*) in day 1 wild-type and *tph-1(ot1274)*.

**(E)** Quantification of PHB>AVA synaptic GRASP (*otIs839*) in wild-type well-fed males supplemented with octopamine (OA) or L1-starved males supplemented with serotonin(5-HT) during the starvation period.

Statistics: (B) Two-way ANOVA followed by Bonferroni multiple comparisons test and (C,D) Student *t*-test. *p*-value and N numbers are indicated on the graph. Scale bar = 10 µm. + indicates the mean value.

**Figure S2. LIN-29A expression in adult male PHB is controlled by sexual and cellular identity together with juvenile experience. Related to Figure 3.**

**(A)** Quantification of the percentage of PHB neurons expressing *lin-29(xe63[gfp::lin-29a])* in wild-type well-fed males supplemented with octopamine (OA) or L1-starved males supplemented with serotonin (5-HT) during the starvation period.

**(B,C)** Longitudinal analysis of *lin-29(ot1482[lin-29::SL2::GFP::H2B])* expression in AVA neuron. Representative images (B) and quantification (C) of expression of AVA neuron expressing *lin-29(ot1482[lin-29::SL2::GFP::H2B])* in different time points after food exposure.
Statistics: *chi*-squared tests followed by Bonferroni multiple comparisons test. *p*-value and N numbers are indicated on the graph. The red dashed circle indicates PHB. Scale bar = 5 μm. + indicates the mean value.

**Figure S3. Time course of *lin-29a* transcription and LIN-29A protein expression Related to Figure 4.**
**(A)** Representative images (left) and quantification (right) of the percentage of PHB neurons expressing *lin-29(ot1482[lin-29::SL2::GFP::H2B])* and (*lin-29(xe63[gfp::lin-29a])* in wild-type well-fed hermaphrodites and males in L1 stage.
**(B)** Representative images (left) and quantification (right) of the percentage of AVA neurons expressing *lin-29(ot1482[lin-29::SL2::GFP::H2B])* and (*lin-29(xe63[gfp::lin-29a])* in wild-type well-fed hermaphrodites and males in L1 stage.
**(C)** Representative images (left) and quantification (right) of the percentage of PHB neurons expressing *lin-29(ot1482[lin-29::SL2::GFP::H2B])* and (*lin-29(xe63[gfp::lin-29a])* in wild-type well-fed hermaphrodites and males in L2 stage.
**(D)** Representative images (left) and quantification (right) of the percentage of AVA neurons expressing *lin-29(ot1482[lin-29::SL2::GFP::H2B])* and (*lin-29(xe63[gfp::lin-29a])* in wild-type well-fed hermaphrodites and males in L2 stage.
**(E)** Representative images (left) and quantification (right) of the percentage of PHB neurons expressing *lin-29(ot1482[lin-29::SL2::GFP::H2B])* and (*lin-29(xe63[gfp::lin-29a])* in wild-type well-fed hermaphrodites and males in L3 stage.
**(F)** Representative images (left) and quantification (right) of the percentage of AVA neurons expressing *lin-29(ot1482[lin-29::SL2::GFP::H2B])* and (*lin-29(xe63[gfp::lin-29a])* in wild-type well-fed hermaphrodites and males in L3 stage.
**(G)** Representative images (left) and quantification (right) of the percentage of PHB neurons expressing *lin-29(ot1482[lin-29::SL2::GFP::H2B])* and (*lin-29(xe63[gfp::lin-29a])* in wild-type well-fed hermaphrodites and males in L4 stage.

**(H)** Representative images (left) and quantification (right) of the percentage of AVA neurons expressing *lin-29(ot1482[lin-29::SL2::GFP::H2B])* and (*lin-29(xe63[gfp::lin-29a])* in wild-type well-fed hermaphrodites and males in L4 stage.

**(I)** Representative images (left) and quantification (right) of the percentage of PHB neurons expressing *lin-29(ot1482[lin-29::SL2::GFP::H2B])* and (*lin-29(xe63[gfp::lin-29a])* in wild-type well-fed hermaphrodites and males in day 1 stage.

**(J)** Representative images (left) and quantification (right) of the percentage of AVA neurons expressing *lin-29(ot1482[lin-29::SL2::GFP::H2B])* and (*lin-29(xe63[gfp::lin-29a])* in wild-type well-fed hermaphrodites and males in day 1 stage.

**(K)** Schematic illustration of the regulation of lin-29a transcription and translation during the development in PHB and AVA, respectively.

**Figure S4. Synapses visualized by juxtaposed PHB-localized CLA-1 and AVA-localized AVR-14 is affected by *lin-29a*. Related to Figure 5.**

**(A)** Representative images of AVA-juxtaposed GFP::CLA-1 in PHB (*otIs883;otEx8040*) in L3 and day 1 wild-type and *lin-29(xe38)*.

**(B)** Quantification of AVA-juxtaposed GFP::CLA-1 in PHB(*otIs883; otEx8040*) in L3 and day 1 wild-type and *lin-29(xe38)*.

**(C)** Representative images of PHB-juxtaposed AVR-14::TagRFP (*otIs902*) in L3 and day 1 wild-type and *lin-29(xe38)*.

**(D)** Quantification of AVR-14::TagRFP in AVA (*otIs902; him-8(e1489)*) in L3 and day 1 wild-type and *lin-29(xe38)*.

**(E)** Representative images of the juxtaposition of PHB-localized GFP::CLA-1 and AVA-localized AVR-14::TagRFP (*otEx8163*) in L3 and day 1 wild-type and *lin-29(xe38)* hermaphrodites (top) and males (bottom).

**(F)**Quantification of juxtaposition index in in L3 and day 1 wild-type and *lin-29(xe38)* hermaphrodites and males.

Statistics: Two-way ANOVA followed by Bonferroni multiple comparisons test. *p*-value and N numbers are indicated on the graph. Scale bar = 10 µm. + indicates the mean value.

**Figure S5. Loss of *lin-29a* does not affect other sexually dimorphic synapses. Related to Figure 5.**

**(A)** Diagram of sexually dimorphic connections with neurons expressing LIN-29A upon sexual maturation.

**(B,C)** Representative images (B) and quantification (C)of ADL>AVA synaptic GRASP *(otEx6829)* in day 1 wild-type, *lin-29(xe38)* and *lin-29(xe40)* in both sexes.

**(D,E)** Representative images (D) and quantification (E) of PHA>AVG iBLINC (*otIs630*) in day 1 wild-type, *lin-29(xe38)* and *lin-29(xe40)* in both sexes.

**(F,G)** Representative images (F) and (G) quantification of PHB>AVG synaptic GRASP (*otIs614*) in day 1 wild-type and *lin-29(xe38)* in both sexes.

Statistics: (C,E,G)Two-way ANOVA followed by Bonferroni multiple comparisons test. *p*-value and N numbers are indicated on the graph. Scale bar = 10 µm. + indicates the mean value.

**Figure S6. Sexual identity acts through *lin-29a* in the PHB to control PHB>AVA sexual dimorphism. Related to Figure 5.**

**(A)** Quantification of PHB>AVA synaptic GRASP(*otIs839*) in wild-type and *lin-29(xe38)* animals with transgene expressing PHB::FEM-3(*otEx7916* and *otEx8164*).

**(B)** Quantification of PHB>AVA synaptic GRASP(*otIs839*) in wild-type and *lin-29(xe38)* animals with transgene expressing AVA::FEM-3 (*otEx8165*).

**(C)** Quantification of PHB>AVA synaptic GRASP(*otIs839*) in wild-type and *lin-29(xe38)* animals with transgene expressing PHB::GOA-1$^{gof}$(*otEx7925* and *otEx8158*).

Statistics: One-way ANOVA followed by Bonferroni multiple comparisons test. *p*-value and N numbers are indicated on the graph. + indicates the mean value.

**Figure S7. *lin-29a* mutants phenotypes are rescued by the human homolog of *lin29a*. Related to Figure 5,6,7.**

**(A,B)** Quantification of PHB>AVA synaptic GRASP(*otIs839*) in and *lin-29(xe38)* males (A) and wild type hermaphrodites (B) with transgene expressing PHB::LIN-29$^{\Delta Zn}$(*otEx7928* and *otEx7929*).

**(C,D)** Quantification of PHB>AVA synaptic GRASP(*otIs839*) in and *lin-29(xe38)* males (A) and wild type hermaphrodites (B) with transgene expressing PHB::ZNF-362(o*tEx7930* and *otEx7931*).

**(E)** (top) Representative images and (bottom) quantification of PHB neuron expressing *dmd-4(ot935)* in *lin-29(xe38)* males with transgenes that express PHB::LIN-29A *(otEx7961 and otEx7964)* and PHB::ZNF362 *(otEx7997 and otEx7998)*

**(F)** Representative images (top) and quantification (bottom) of *fmi-1(syb4563)* expression in the PHB *lin-29(xe38)* males with transgenes that express PHB::LIN-29A *(otEx7961)* and PHB::ZNF362 *(otEx7997)*

(A,B,C,D) Two-way ANOVA and (E,F) Two-proportion Z test followed by Bonferroni multiple comparisons test. *p*-value and N numbers are indicated on the graph. Scale bar = 5 µm. + indicates the mean value.

**Figure S8. Sexually dimorphic FMI-1 protein expression in the PHB axon is regulated by *lin-29a*. Related to Figure 7.**

**(A)** Schematic illustration for *fmi-1(ot1349)* and *fmi-1(ot1429)*. *fmi-1(ot1349)* is a *fmi-1* allele in which the entire *fmi-1* locus is flanked with CRISPR/Cas9-engineered LoxP sites and a GFP::H2B tagged at the C-terminus region after second LoxP, after the *fmi-1* stop codon of *fmi-1*. Upon excision of *fmi-1* with Cre, the GFP::H2B is then expressed in the cell expressing Cre recombinase. *fmi-1(ot1429)* inserts codon-optimized 6XGFP$_{11}$ at the C-terminus.

**(B)** Schematic illustration of the visualization of the endogenous FMI-1 protein. The other half of GFP with myristoylation peptide (myri-GPF1-10) is expressed in the PHB by a transgene (*otEx8148*) to visualize the membrane-localized FMI-1.

**(C,D)** Representative images (C) and quantification (D) of endogenous FMI-1 expression from *fmi-1(ot1429); otEx8148* in day 1 wild type and *lin-29(xe38)* of both sexes.

Two-way ANOVA followed by Bonferroni multiple comparisons test. *p*-value and N numbers are indicated on the graph. Scale bar = 10 µm. + indicates the mean value.

**Figure S9. Loss of *fmi-1* results in aberrant PHB>AVA connectivity. Related to Figure 8.**

**(A)** Representative images (top) and quantification (bottom) of AVA-juxtaposed GFP::CLA-1 in PHB (*otIs883;otEx8040*) in day 1 wild-type and *fmi-1(ot1090)* hermaphrodites. *fmi-1(ot1090)* is also the *fmi-1* locus deletion, which removes all *fmi-1* isoforms, similar to that of *fmi-1(ot1291)*.

**(B)** Representative images (top) and quantification (D) of PHB-juxtaposed AVR-14::TagRFP (*otIs902*) in day 1 wild-type and *fmi-1(ot1090)* hermaphrodites.

**(C)** Representative images of the juxtaposition of PHB-localized GFP::CLA-1 and AVA-localized AVR-14::TagRFP (*otEx8163*) in day 1 wild-type and *fmi-1(ot1291)* hermaphrodites.

**(D)** Quantification of the juxtaposition of PHB-localized GFP::CLA-1 and AVA-localized AVR-14::TagRFP (*otEx8163*) in day 1 wild-type and *fmi-1(ot1291)* hermaphrodites.

(A,B,D) Student *t*-test and. *p*-value and N numbers are indicated on the graph. Scale bar = 10 μm. + indicates the mean value.

**Figure S10. Schematic summary of the study.**

**(A)** LIN-29A integrates four dimensions of information, including cell identity (CEH-14), sexual identity (TRA-1), temporal identity (LIN-41), and feeding state experience (Serotonin-SER-4-CRH-1).

**(B)** Schematic illustration of FMI-1 functions in PHB embryonic and postembryonic developmental stages and adulthood.

**(C)** Schematic illustration of the regulation of LIN-29A>DMD-4>*fmi-1* in the nucleus and the corresponding regulation of PHB>AVA neurite adjacency and synaptogenesis.

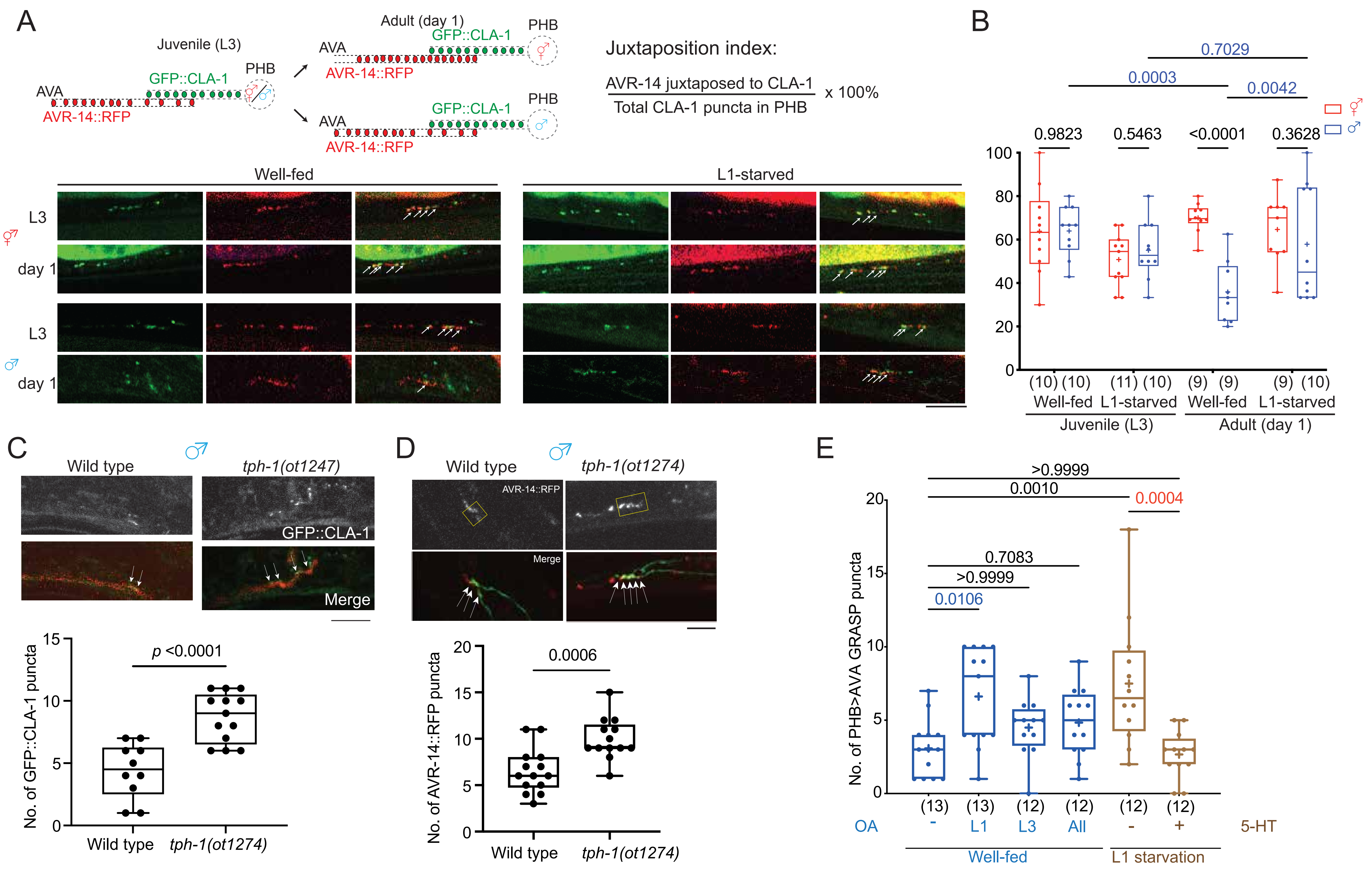

A
Juvenile (L3)
Adult (day 1)
AVA
PHB
GFP::CLA-1
AVR-14::RFP
Juxtaposition index:
AVR-14 juxtaposed to CLA-1
Total CLA-1 puncta in PHB
x 100%
Well-fed
L1-starved
L3
day 1
B
0.7029
0.0003
0.0042
0.9823
0.5463
<0.0001
0.3628
(10) (10)
(11) (10)
(9) (9)
(9) (10)
Well-fed
L1-starved
Well-fed
L1-starved
Juvenile (L3)
Adult (day 1)
C
Wild type
tph-1(ot1247)
GFP::CLA-1
Merge
p <0.0001
No. of GFP::CLA-1 puncta
Wild type
tph-1(ot1274)
D
Wild type
tph-1(ot1274)
AVR-14::RFP
Merge
0.0006
No. of AVR-14::RFP puncta
Wild type
tph-1(ot1274)
E
>0.9999
0.0010
0.0004
0.7083
>0.9999
0.0106
No. of PHB>AVA GRASP puncta
(13)
(13)
(12)
(12)
(12)
(12)
OA
-
L1
L3
All
-
+
5-HT
Well-fed
L1 starvation


Fig S1

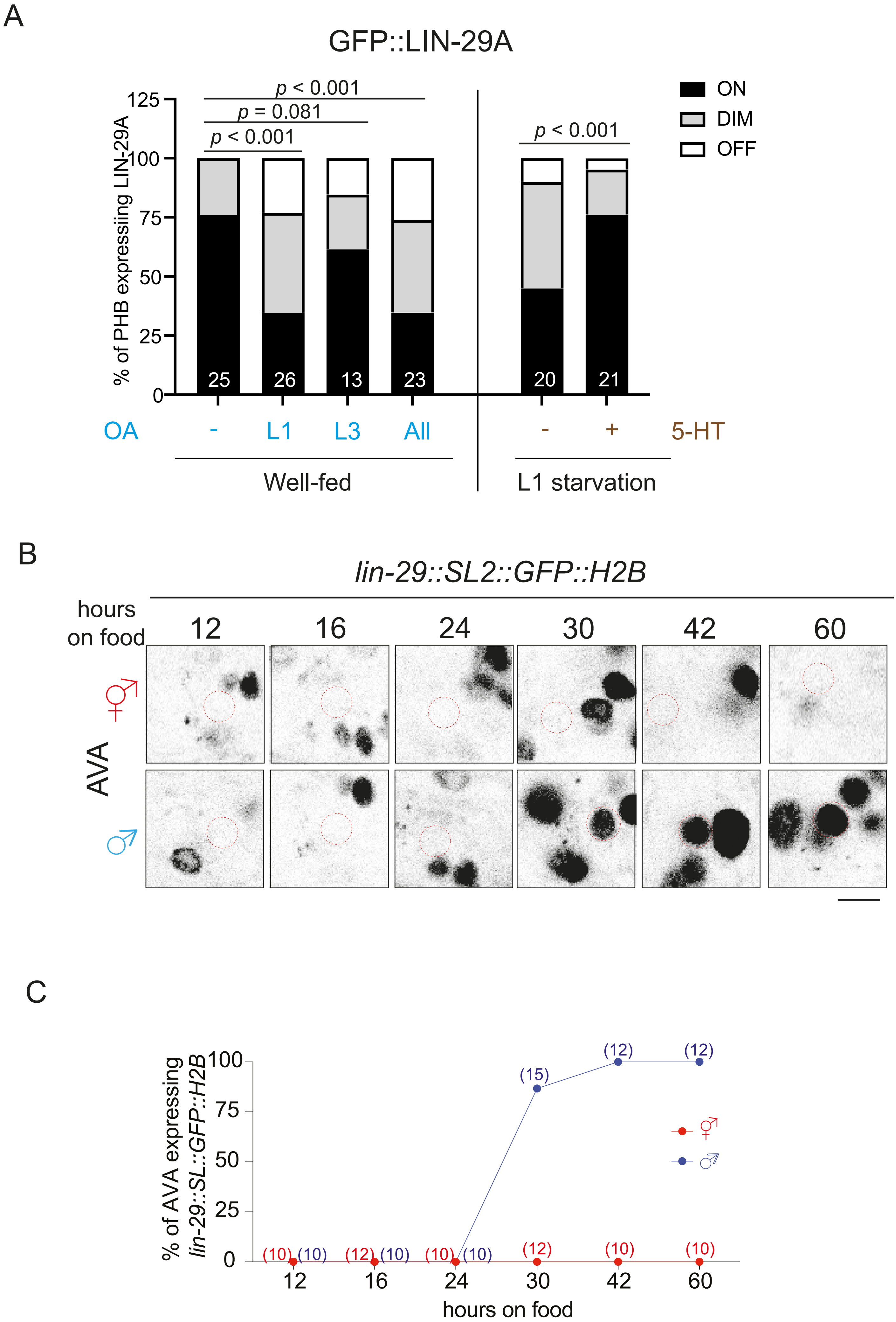

A
GFP::LIN-29A
% of PHB expressing LIN-29A
125
100
75
50
25
0
p < 0.001
p = 0.081
p < 0.001
p < 0.001
ON
DIM
OFF
25
26
13
23
20
21
OA
-
L1
L3
All
-
+
5-HT
Well-fed
L1 starvation
B
lin-29::SL2::GFP::H2B
hours on food
12
16
24
30
42
60
AVA
♀
♂
C
% of AVA expressing lin-29::SL::GFP::H2B
100
75
50
25
0
(15)
(12)
(12)
(10)
(10)
(12)
(10)
(10)
(10)
(12)
(10)
(10)
12
16
24
30
42
60
hours on food


Fig S2

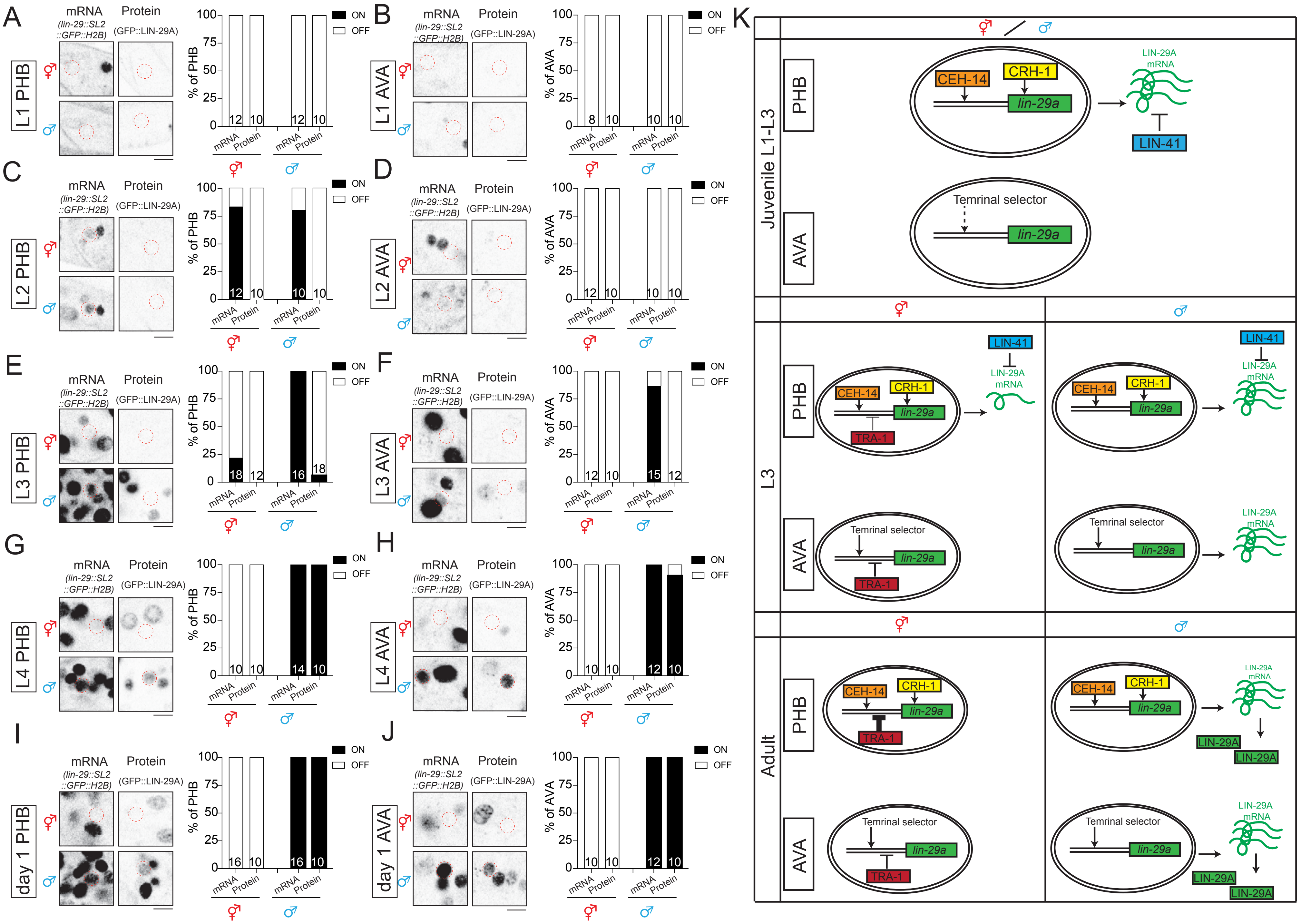
A
L1 PHB
B
L1 AVA
C
L2 PHB
D
L2 AVA
E
L3 PHB
F
L3 AVA
G
L4 PHB
H
L4 AVA
I
day 1 PHB
J
day 1 AVA
mRNA
(lin-29::SL2::GFP::H2B)
Protein
(GFP::LIN-29A)
% of PHB
% of AVA
ON
OFF
K
Juvenile L1-L3
L3
Adult
PHB
AVA
CEH-14
CRH-1
lin-29a
LIN-29A mRNA
LIN-41
Temrinal selector
TRA-1
LIN-29A

Fig S3

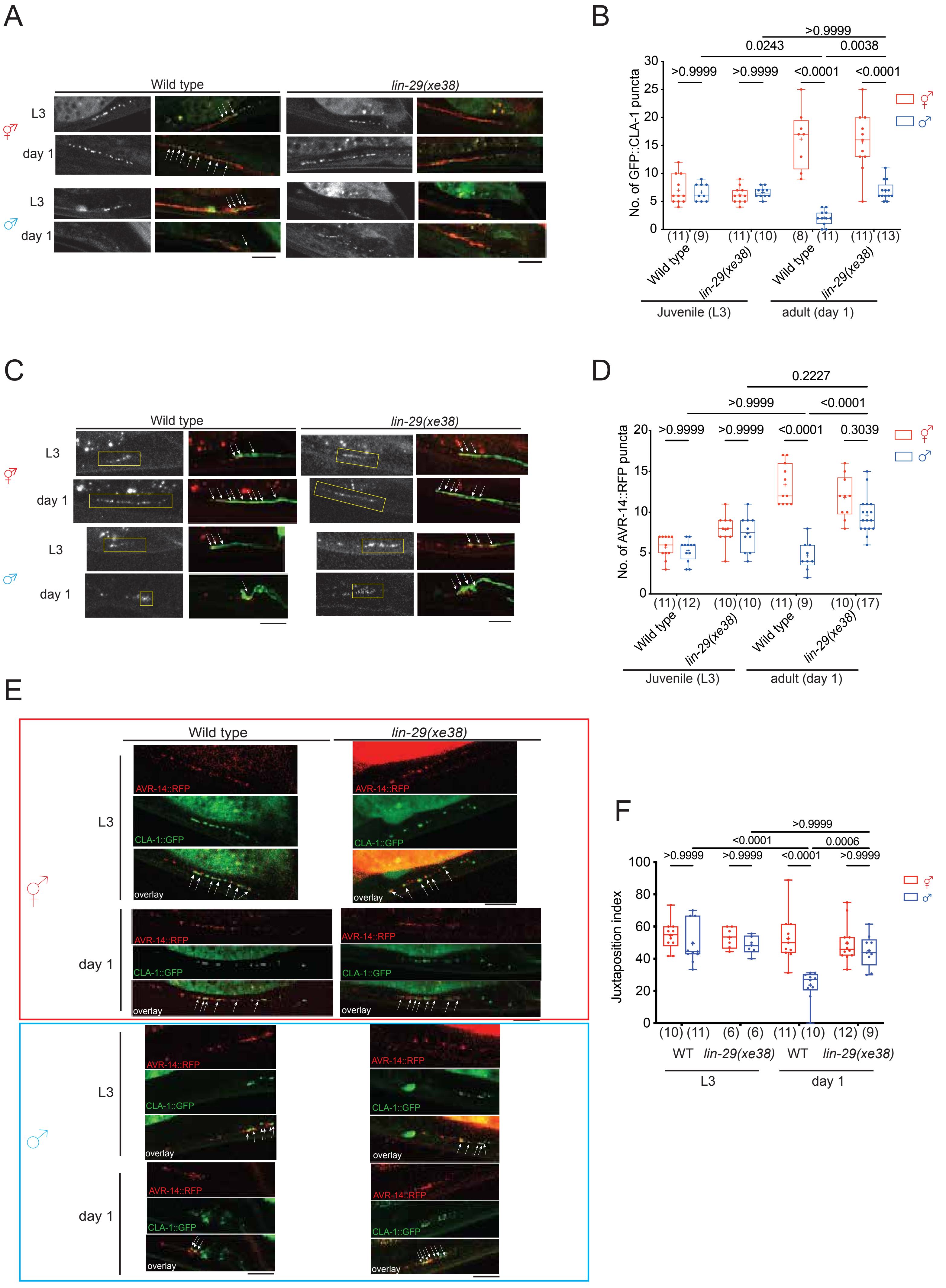

A
Wild type
lin-29(xe38)
L3
day 1
B
No. of GFP::CLA-1 puncta
>0.9999
0.0243
0.0038
<0.0001
(11) (9) (11) (10) (8) (11) (11) (13)
Wild type
lin-29(xe38)
Juvenile (L3)
adult (day 1)
C
D
No. of AVR-14::RFP puncta
0.2227
0.3039
(11) (12) (10) (10) (11) (9) (10) (17)
E
AVR-14::RFP
CLA-1::GFP
overlay
F
Juxtaposition index
0.0006
(10) (11) (6) (6) (11) (10) (12) (9)
WT


Fig S4

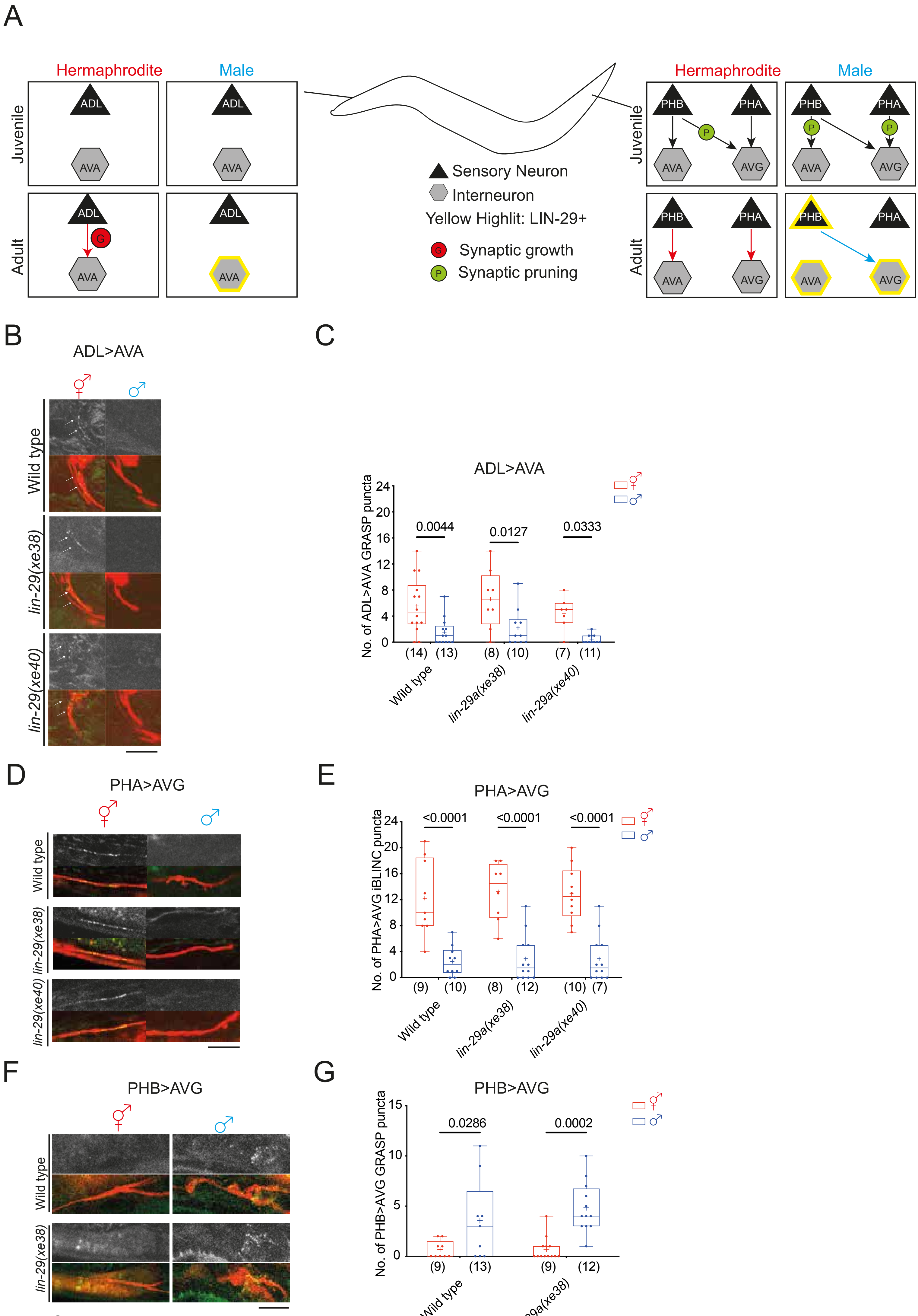
A
Hermaphrodite
Male
Juvenile
Adult
ADL
AVA
G
PHB
PHA
AVG
P
Sensory Neuron
Interneuron
Yellow Highlit: LIN-29+
Synaptic growth
Synaptic pruning
B
ADL>AVA
Wild type
lin-29(xe38)
lin-29(xe40)
C
ADL>AVA
No. of ADL>AVA GRASP puncta
0.0044
0.0127
0.0333
(14) (13)
(8) (10)
(7) (11)
Wild type
lin-29a(xe38)
lin-29a(xe40)
D
PHA>AVG
Wild type
lin-29(xe38)
lin-29(xe40)
E
PHA>AVG
No. of PHA>AVG iBLINC puncta
<0.0001
<0.0001
<0.0001
(9) (10)
(8) (12)
(10) (7)
Wild type
lin-29a(xe38)
lin-29a(xe40)
F
PHB>AVG
Wild type
lin-29(xe38)
G
PHB>AVG
No. of PHB>AVG GRASP puncta
0.0286
0.0002
(9) (13)
(9) (12)
Wild type
lin-29a(xe38)

Fig S5

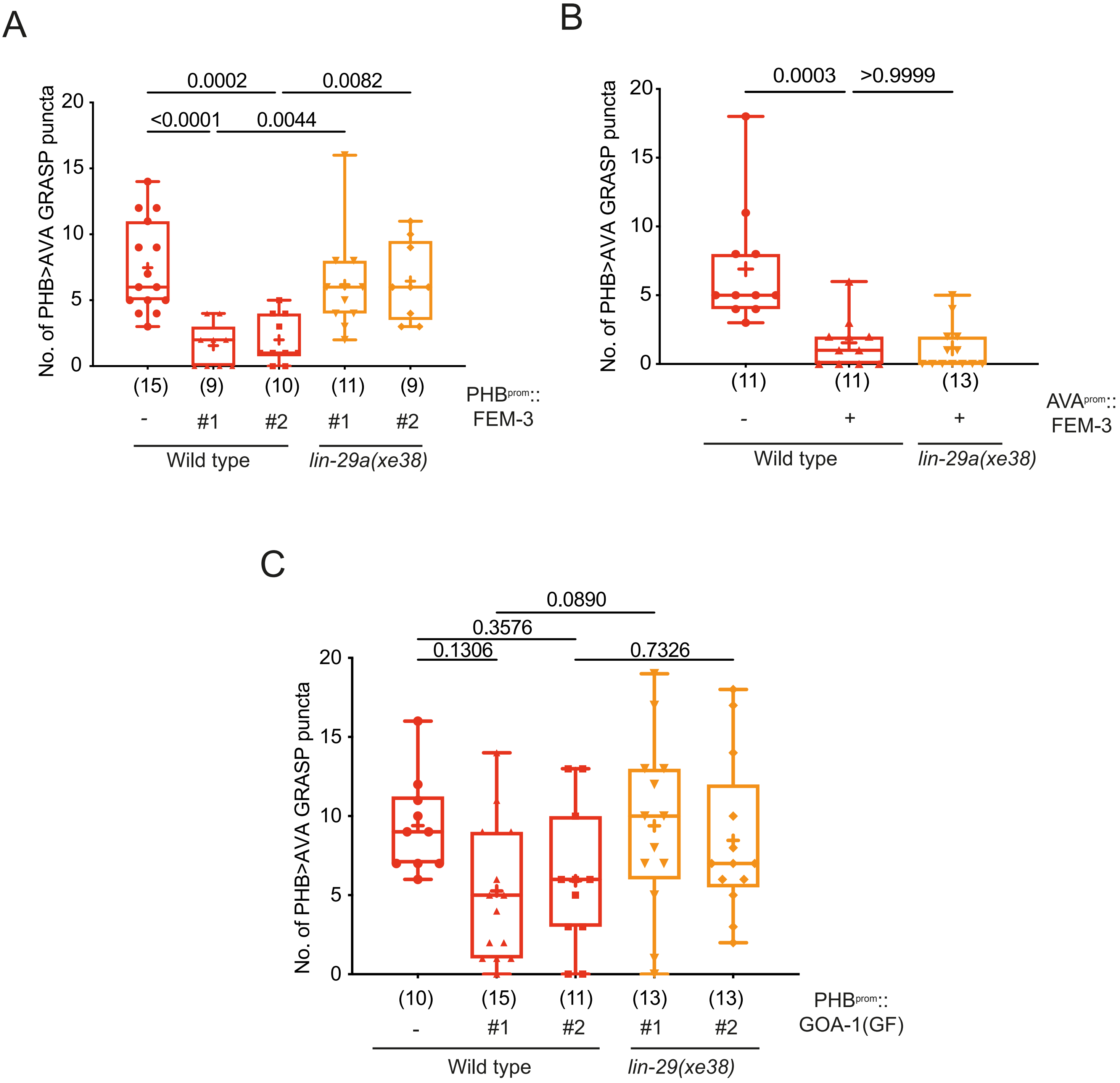
A
0.0002
0.0082
<0.0001
0.0044
No. of PHB>AVA GRASP puncta
20
15
10
5
0
(15)
(9)
(10)
(11)
(9)
PHBprom::
FEM-3
-
#1
#2
#1
#2
Wild type
lin-29a(xe38)
B
0.0003
>0.9999
No. of PHB>AVA GRASP puncta
20
15
10
5
0
(11)
(11)
(13)
AVAprom::
FEM-3
-
+
+
Wild type
lin-29a(xe38)
C
0.0890
0.3576
0.1306
0.7326
No. of PHB>AVA GRASP puncta
20
15
10
5
0
(10)
(15)
(11)
(13)
(13)
PHBprom::
GOA-1(GF)
-
#1
#2
#1
#2
Wild type
lin-29(xe38)

Fig S6

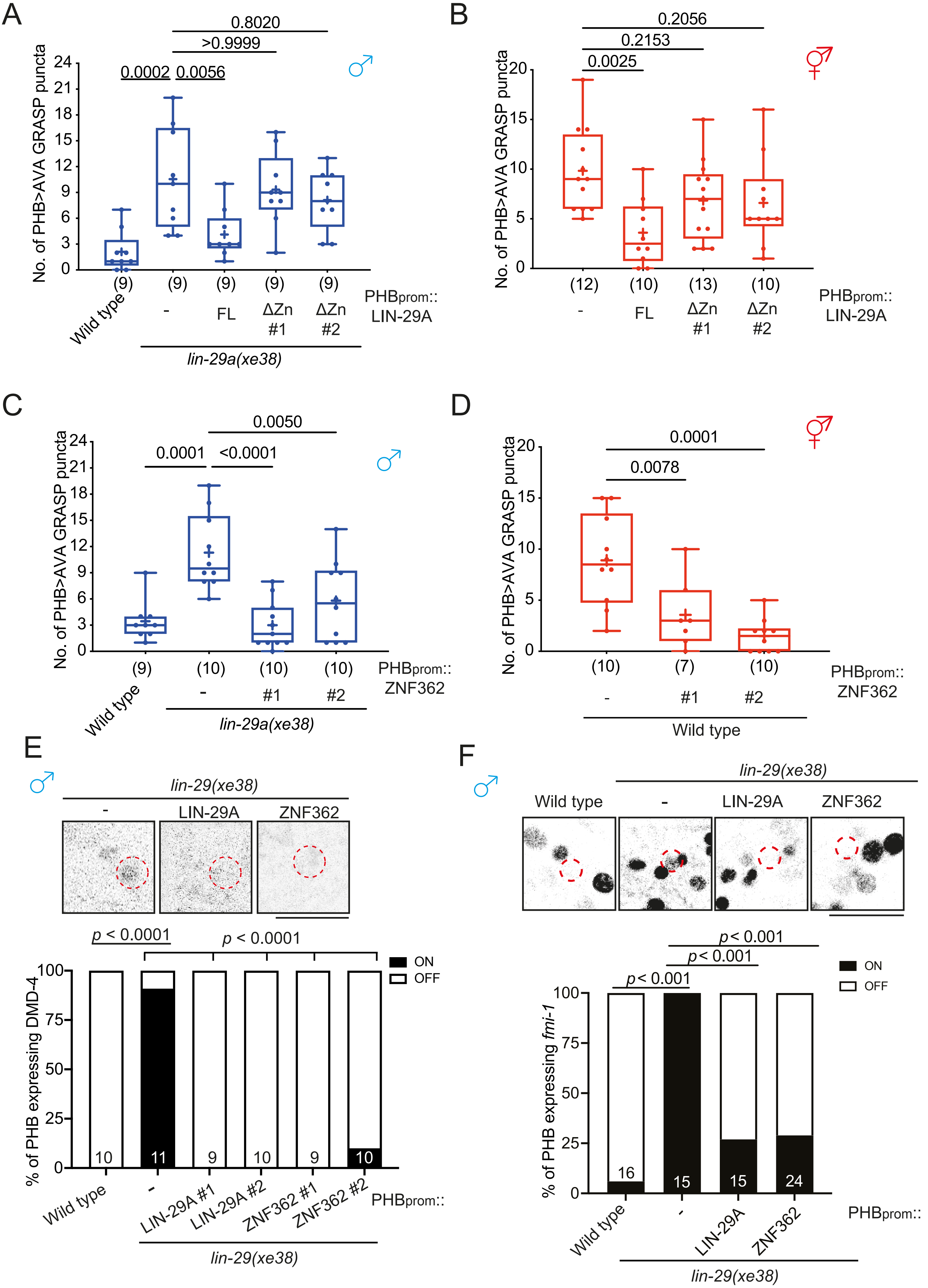
A
B
C
D
E
F
No. of PHB>AVA GRASP puncta
0.8020
>0.9999
0.0002
0.0056
Wild type
FL
ΔZn #1
ΔZn #2
PHBprom:: LIN-29A
lin-29a(xe38)
0.2056
0.2153
0.0025
0.0050
0.0001
<0.0001
PHBprom:: ZNF362
0.0078
lin-29(xe38)
LIN-29A
ZNF362
p < 0.0001
p < 0.001
ON
OFF
% of PHB expressing DMD-4
% of PHB expressing fmi-1
LIN-29A #1
LIN-29A #2
ZNF362 #1
ZNF362 #2
PHBprom::

Fig S7

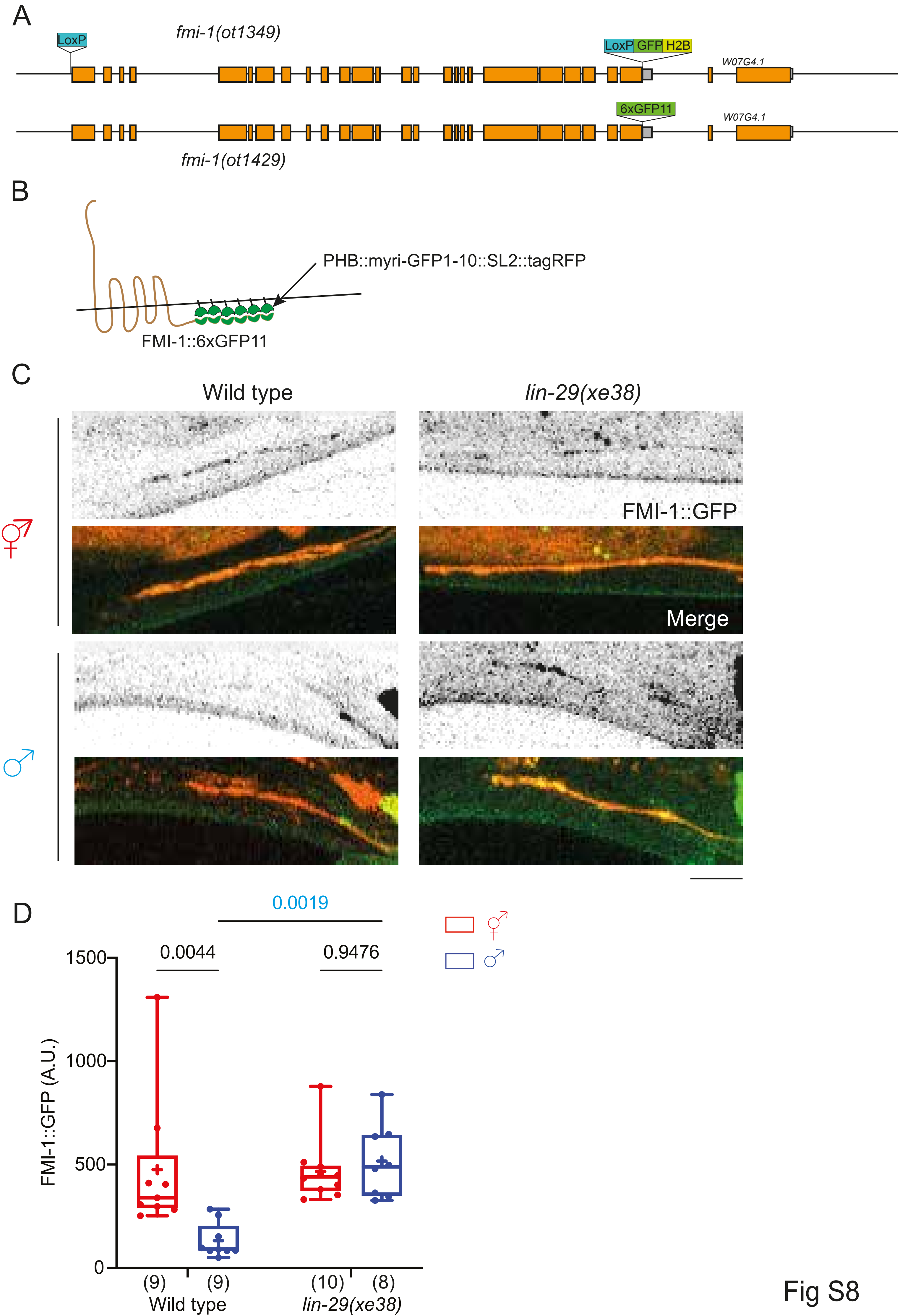
A
LoxP
fmi-1(ot1349)
LoxP GFP H2B
W07G4.1
6xGFP11
W07G4.1
fmi-1(ot1429)
B
PHB::myri-GFP1-10::SL2::tagRFP
FMI-1::6xGFP11
C
Wild type
lin-29(xe38)
FMI-1::GFP
Merge
D
0.0019
0.0044
0.9476
1500
1000
500
0
FMI-1::GFP (A.U.)
(9)
(9)
(10)
(8)
Wild type
lin-29(xe38)

Fig S8

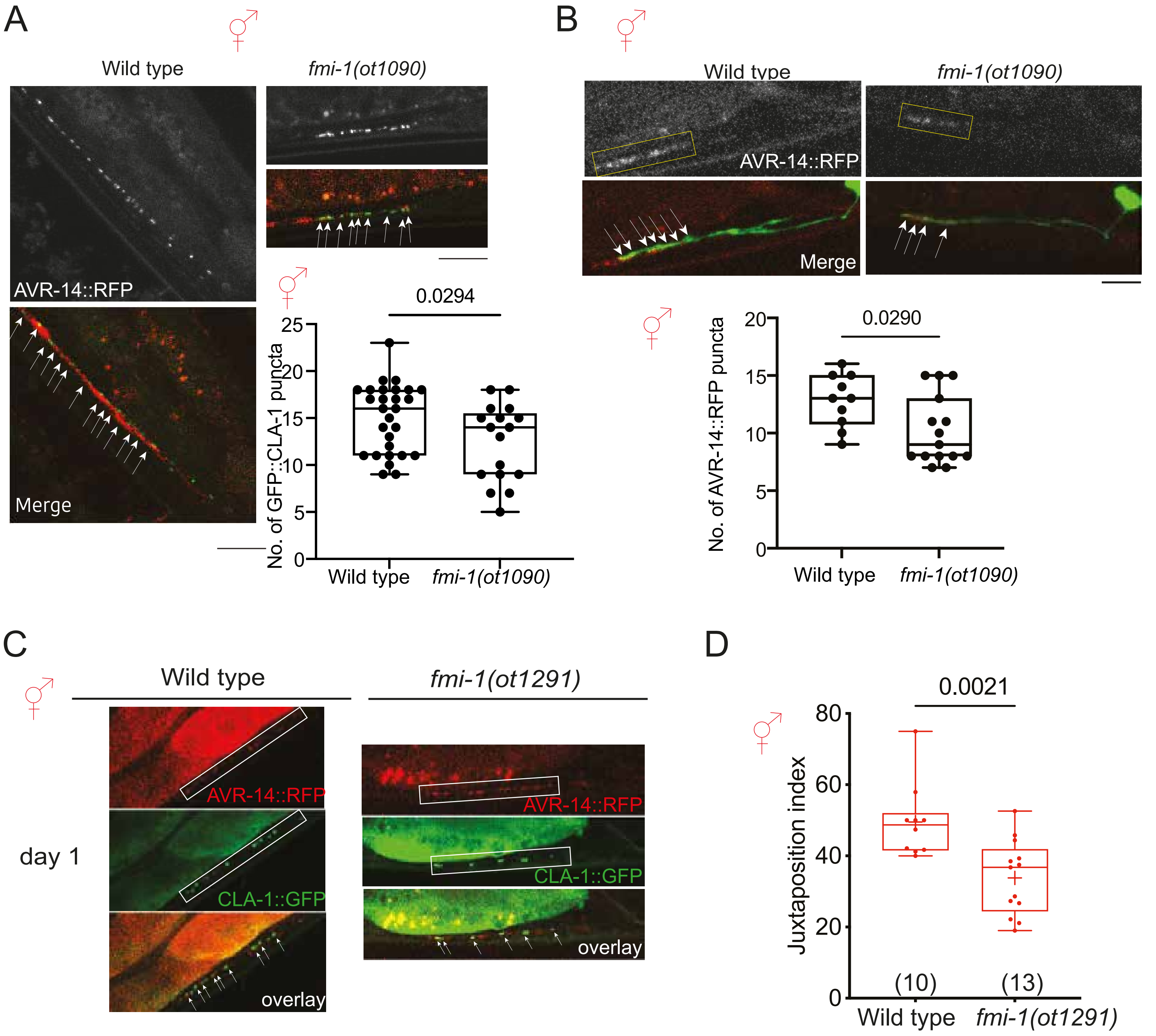
A
Wild type
fmi-1(ot1090)
AVR-14::RFP
Merge
0.0294
No. of GFP::CLA-1 puncta
25
20
15
10
5
0
Wild type
fmi-1(ot1090)
B
Wild type
fmi-1(ot1090)
AVR-14::RFP
Merge
0.0290
No. of AVR-14::RFP puncta
20
15
10
5
0
Wild type
fmi-1(ot1090)
C
Wild type
fmi-1(ot1291)
day 1
AVR-14::RFP
CLA-1::GFP
overlay
AVR-14::RFP
CLA-1::GFP
overlay
D
0.0021
Juxtaposition index
80
60
40
20
0
(10)
(13)
Wild type
fmi-1(ot1291)

Fig S9

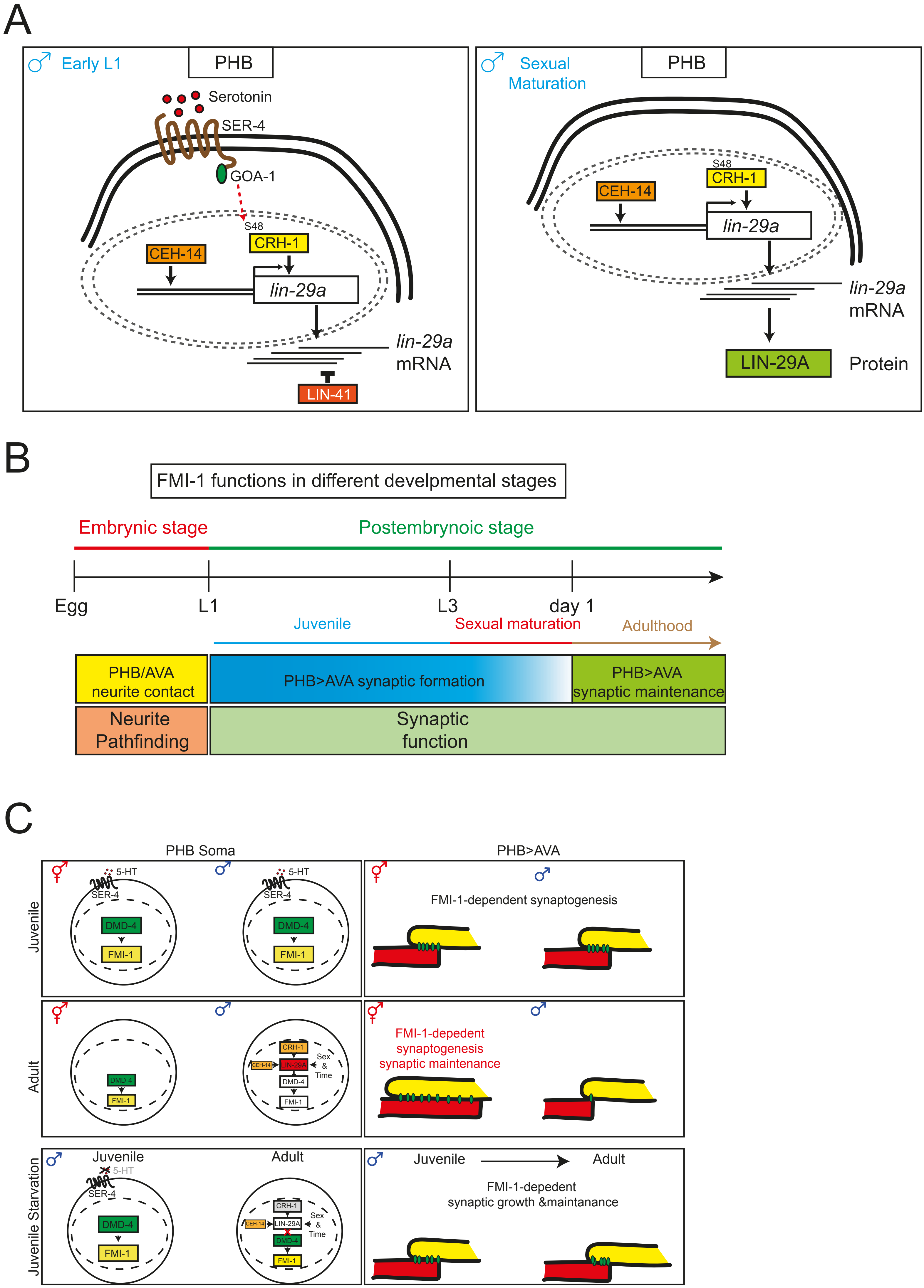
A
Early L1
PHB
Serotonin
SER-4
GOA-1
S48
CRH-1
CEH-14
lin-29a
lin-29a mRNA
LIN-41
Sexual Maturation
PHB
S48
CRH-1
CEH-14
lin-29a
lin-29a mRNA
LIN-29A
Protein
B
FMI-1 functions in different develpmental stages
Embrynic stage
Postembrynoic stage
Egg
L1
L3
day 1
Juvenile
Sexual maturation
Adulthood
PHB/AVA neurite contact
PHB>AVA synaptic formation
PHB>AVA synaptic maintenance
Neurite Pathfinding
Synaptic function
C
PHB Soma
PHB>AVA
Juvenile
5-HT
SER-4
DMD-4
FMI-1
FMI-1-dependent synaptogenesis
Adult
CRH-1
CEH-14
LIN-29A
Sex & Time
FMI-1-depedent synaptogenesis synaptic maintenance
Juvenile Starvation
Juvenile
Adult
FMI-1-depedent synaptic growth &maintanance

Fig S10
